# Dopamine D1 receptor activation drives plasticity in the songbird auditory pallium

**DOI:** 10.1101/2020.10.09.330266

**Authors:** Matheus Macedo-Lima, Hannah M. Boyd, Luke Remage-Healey

**Author notes:** Competing interests: The authors declare no competing interests.

## Abstract

Vocal learning species must form and extensively hone associations between sounds and social contingencies. In songbirds, dopamine signaling guides song motor-production, variability, and motivation, but it is unclear how dopamine regulates fundamental auditory associations for learning new sounds. We hypothesized that dopamine regulates learning in the auditory pallium, in part by interacting with local neuroestradiol signaling. Here, we show that zebra finch auditory neurons frequently coexpress D1 receptor (D1R) protein, neuroestradiol-synthase, GABA, and parvalbumin. Auditory classical conditioning increased neuroplasticity gene induction in D1R-positive neurons. *In vitro,* D1R pharmacological activation reduced the amplitude of GABAergic and glutamatergic currents, and increased the latter’s frequency. *In vivo*, D1R activation reduced the firing of putative interneurons, increased the firing of putative excitatory neurons, and made both neuronal types unable to adapt to novel stimuli. Together, these data support the hypothesis that dopamine acting via D1Rs modulates learning and memory in the songbird sensory cortex.

## 1.1 Introduction

Vocal and auditory learning have evolved in select few organisms, including humans and songbirds. Studying vocal learning in songbirds continues to provide insight into the suite of mechanisms that support spoken language (Jarvis, 2019). The first step in the spoken language learning process is to make socially-contingent associations about complex sounds. This likely engages cortical brain structures, where sounds and meaning are bound.

Midbrain structures that process reinforcement can guide the formation of auditory associations in the cortex through the release of the neuromodulator dopamine. In mammals, dopamine in the primary auditory cortex drives auditory learning and plasticity of simple stimuli, such as pure tones (Bao et al., 2001; Schicknick et al., 2012, 2008). The role of dopamine and dopaminergic midbrain nuclei have also been examined in the songbird brain, but primarily in the context of reinforcement learning for song production and motivation to sing (Gadagkar et al., 2016; Leblois et al., 2010; Matheson and Sakata, 2015; Schmidt and Ding, 2014; Tanaka et al., 2018). Much less is known about how dopamine signaling may regulate auditory processing and association of complex sounds such as song.

The songbird caudomedial nidopallium (NCM; Fig. 1A) is a pallial secondary auditory region considered analogous to the center for speech comprehension in humans, Wernicke’s area (Bolhuis et al., 2010). Neuronal responses in awake, restrained zebra finch NCM show stimulus-specific adaptation to sounds played repeatedly, consistent with active memory formation (Chew et al., 1996; Lu and Vicario, 2017). This adaptation in NCM neurons reflects familiarity with songs, as well as song-consequence associations (Bell et al., 2015; Chew et al., 1996; Lu and Vicario, 2017). The NCM is an important target of a wide variety of neuromodulators, such as norepinephrine (Ikeda et al., 2015; Lee et al., 2018), nitric oxide (Wallhäusser-Franke et al., 1995), neuroestradiol (Macedo-Lima and Remage-Healey, 2020; Saldanha et al., 2000) and dopamine (Matragrano et al., 2012). In addition to neuromodulators, NCM is rich in NMDA receptors (Saldanha et al., 2004), which are classically regarded as key players in cellular memory formation processes, such as long-term potentiation and long-term depression (Lüscher and Malenka, 2012). NCM therefore is a highly plastic structure involved in the processing, and perhaps association, of complex sounds. However, the molecular and neuromodulatory mechanisms underlying associations between sound and context in higher-order circuits like NCM have not been elucidated.

**Figure 1.**
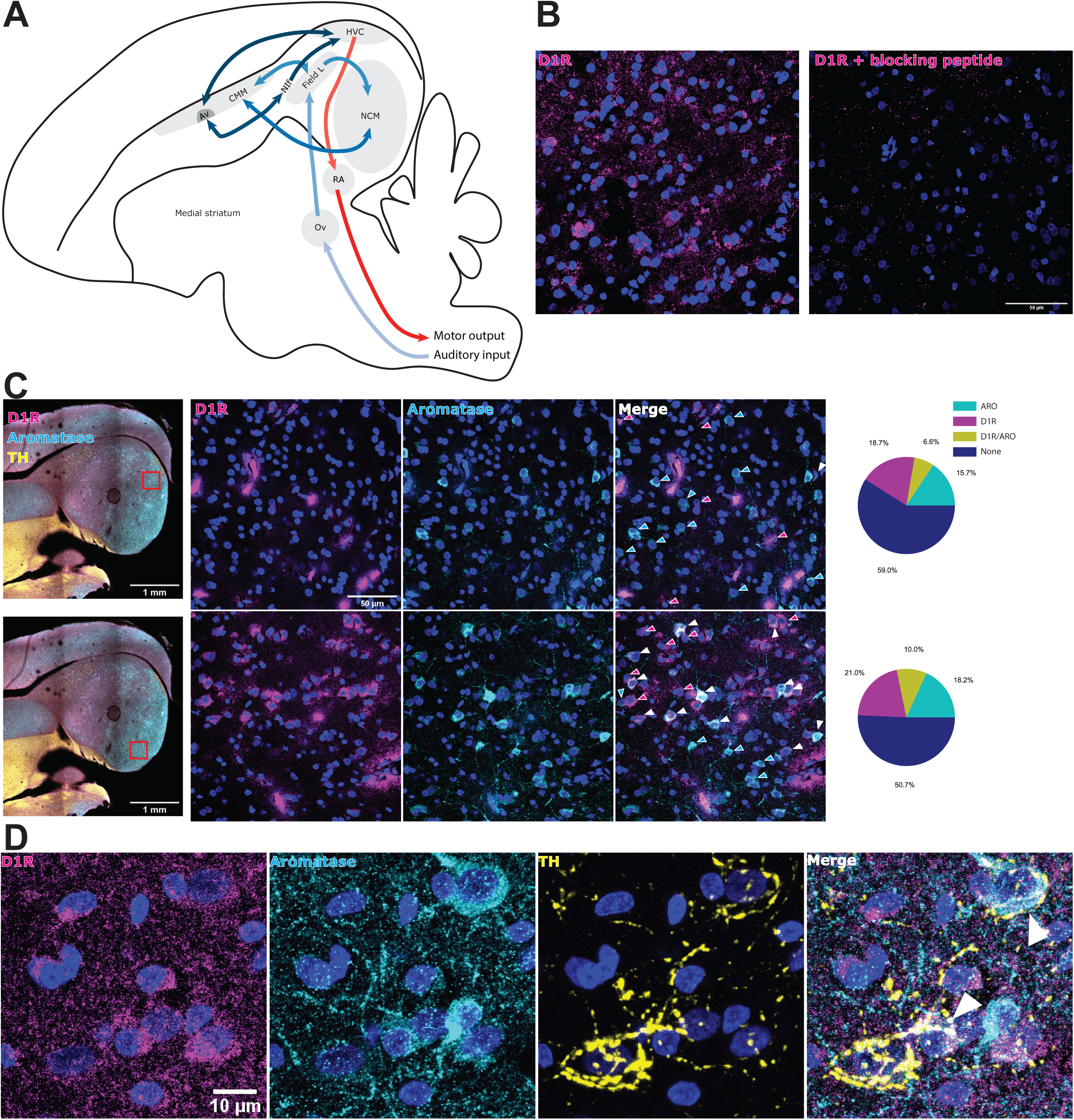
Aromatase and D1 receptor (D1R) proteins are coexpressed by caudomedial nidopallium (NCM) neurons. (A) Songbird auditory circuits. Lighter-to-darker colors illustrate the trajectory of auditory (blue) and motor (red) information. (B) Specificity confirmation of the anti-D1R antibody used here. Preincubation with blocking peptide virtually eliminated cellular staining. (C) Triple immunofluorescence staining for D1R, aromatase and tyrosine hydroxylase (TH; enzyme participating in dopamine synthesis). Magenta and cyan arrows indicate single-labeled D1R and aromatase+ neurons, respectively. White arrows indicate double-labeling. (D) TH fibers are frequently found in close association with double-labeled neurons. Note that fibers tightly envelop double-labeled soma (arrows). Av: nucleus avalanche; ARO: aromatase; CMM: caudomedial mesopallium; D1R: dopamine D1 receptor; HVC: HVC (proper name); Ov: nucleus ovoidallis; NCM: caudomedial nidopallium; NIf: nucleus interfacialis; RA: robust nucleus of the arcopallium; TH: tyrosine hydroxylase.

Dopaminergic fibers permeate the secondary auditory regions NCM and the caudomedial mesopallium (CMM), but not the thalamo-recipient auditory region, Field L (Reiner et al., 1994; von Eugen et al., 2020). Dopamine receptor mRNA maps onto this architecture, such that D1 receptors (D1Rs) are abundant in NCM and CMM, and D2 receptors are abundant in CMM but not in NCM. Neither receptor is evident in the primary auditory pallium Field L (Kubikova et al., 2010). Neurons in the ventral tegmental area have been reported to project to the NCM and could provide a learning-contingent source of dopamine to this region (Yanagihara et al., 2019). Indeed, hearing song increases rapid turnover of dopamine in songbird auditory regions, especially in the NCM (Matragrano et al., 2012). Therefore, dopamine signaling is a feature distinguishing secondary from primary auditory pallium in songbirds, providing an anatomical circuit-basis for dopamine-dependent auditory learning. The extent to which dopamine signaling modulates the encoding and processing of stimuli in songbird neurons, however, is unclear.

In this study, we hypothesized that dopamine signaling via D1 receptors (D1Rs) mediates NCM neural circuit plasticity. We obtained anatomical, behavioral, and physiological evidence that support this hypothesis. First, we used immunofluorescence to characterize D1R-expressing neurons regarding aromatase, GABA, and parvalbumin expression. To ask if D1Rs were involved in auditory learning, we examined the expression of the immediate-early gene EGR1 in D1R-expressing neurons during an auditory classical conditioning task. To characterize the synaptic effects of D1R activation we employed *in vitro* intracellular recordings and recorded glutamate- and GABAergic currents while delivering a D1R agonist. Finally, to examine the functional effects of D1R activation on auditory physiology and plasticity we delivered the D1R agonist during *in vivo* awake extracellular multielectrode recordings and evaluated whether D1R activation modulated neuronal firing patterns and stimulus-specific adaptation in response to auditory signals.

## 1.2 Results

### 1.2.1 Aromatase and D1-receptor proteins are coexpressed by NCM neurons

NCM is a key region for auditory processing in songbirds (Fig. 1A). Previously, we showed that neuroestradiol production in NCM participated in reinforcement-driven auditory association learning (Macedo-Lima and Remage-Healey, 2020). Thus, we hypothesized that reinforcement signaling by the dopaminergic system involved interactions with aromatase neurons in NCM. D1 receptor (D1R) mRNA has been previously reported in NCM (Kubikova et al., 2010), but not its end product, D1R protein. Here we identified a working antibody for the specific detection of D1R protein in songbirds as confirmed by the absence of staining when tissue was preincubated with the D1R antigen (Fig. 1B). We found that D1R+ neurons were often found coexpressing aromatase, representing 29.6 and 35.4% of the aromatase+ (ARO+) neuronal subpopulation in dNCM and vNCM respectively (Fig. 1C). Moreover, the population of D1R+/ARO+ neurons represented 6.6 and 10% of the neuronal population (DAPI nuclei) in dNCM and vNCM, respectively. A chi-square test analyzing these proportions yielded a significant relationship between D1R and ARO counts (χ^2^(1)=10.926, p<0.001), with double-labeled cells occurring significantly more frequently than expected (ARO+/D1R+ Pearson’s residual=2.476, p=0.007). Interestingly, we frequently observed tyrosine hydroxylase+ (TH+) fibers enveloping D1R+/ARO+ neurons (see Fig. 1D for representative images). We report anatomical distributions of D1R/ARO expression in the supplementary material (Supplementary Fig. 1). These data show that D1R protein is prevalent in NCM neurons and suggest that D1R+ neurons represent a significant population in NCM. Moreover, D1R+ neurons are significantly more likely to express ARO+ than D1R– neurons, suggesting an association between estrogen production and dopaminergic signaling.

### 1.2.2 Song induction of EGR1 in D1R+ and ARO+ neurons

Next, we examined whether D1R+ and ARO+ NCM neurons were involved in processing of conspecific songs, using the neuroplasticity marker EGR1. Conspecific song playback to birds significantly increased overall EGR1 expression in NCM when compared to birds that experienced silence (Fig. 2A-B; Supplementary Fig. 2B), which is well established in the literature (Mello and Clayton, 1994; Mello et al., 1992). GLM/ANOVA showed that the degree of song induction of neuronal EGR1 (% of NeuN) was stronger in L-vNCM and R-dNCM, but consistently higher on average across all areas (Supplementary Fig. 2B; Hemisphere*Area*Condition: χ^2^(1)=7.748, p=0.005; Tukey’s post-hoc test Song – Silence: L-vNCM: t(18)=2.887, p=0.010, L-dNCM: t(18)=1.820, p=0.085, R-vNCM: t(18)=1.180, p=0.253, R-dNCM: t(18)=5.012, p<0.001). We next showed that EGR1 song induction was also increased in D1R+ neurons (Fig. 2B; Area (dNCM>vNCM): χ^2^(1)=10.202, p=0.001; Condition: χ^2^(1)=6.774, p=0.009; p>0.08 in all other main effects), in ARO+ neurons (Fig. 2B; Area (dNCM>vNCM): χ^2^(1)=56.723, p<0.001; Condition: χ^2^(1)=13.824, p<0.001; p>0.061 in all other main effects) and in double labeled D1R+/ARO+ neurons (Fig. 2B; Area (dNCM>vNCM):χ^2^(1)=3.978, p=0.046; Condition: χ^2^(1)=4.491, p=0.034; Area*Condition: χ^2^(1)=3.613, p=0.057; all other p>0.190). These data show that song stimulation increased EGR1 expression in D1R+, ARO+ and double-labeled D1R+/ARO+ neurons. These results are consistent with the notion that dopamine and neuroestrogen signaling are each involved in auditory responses to song, and may interact in that capacity.

**Figure 2.**
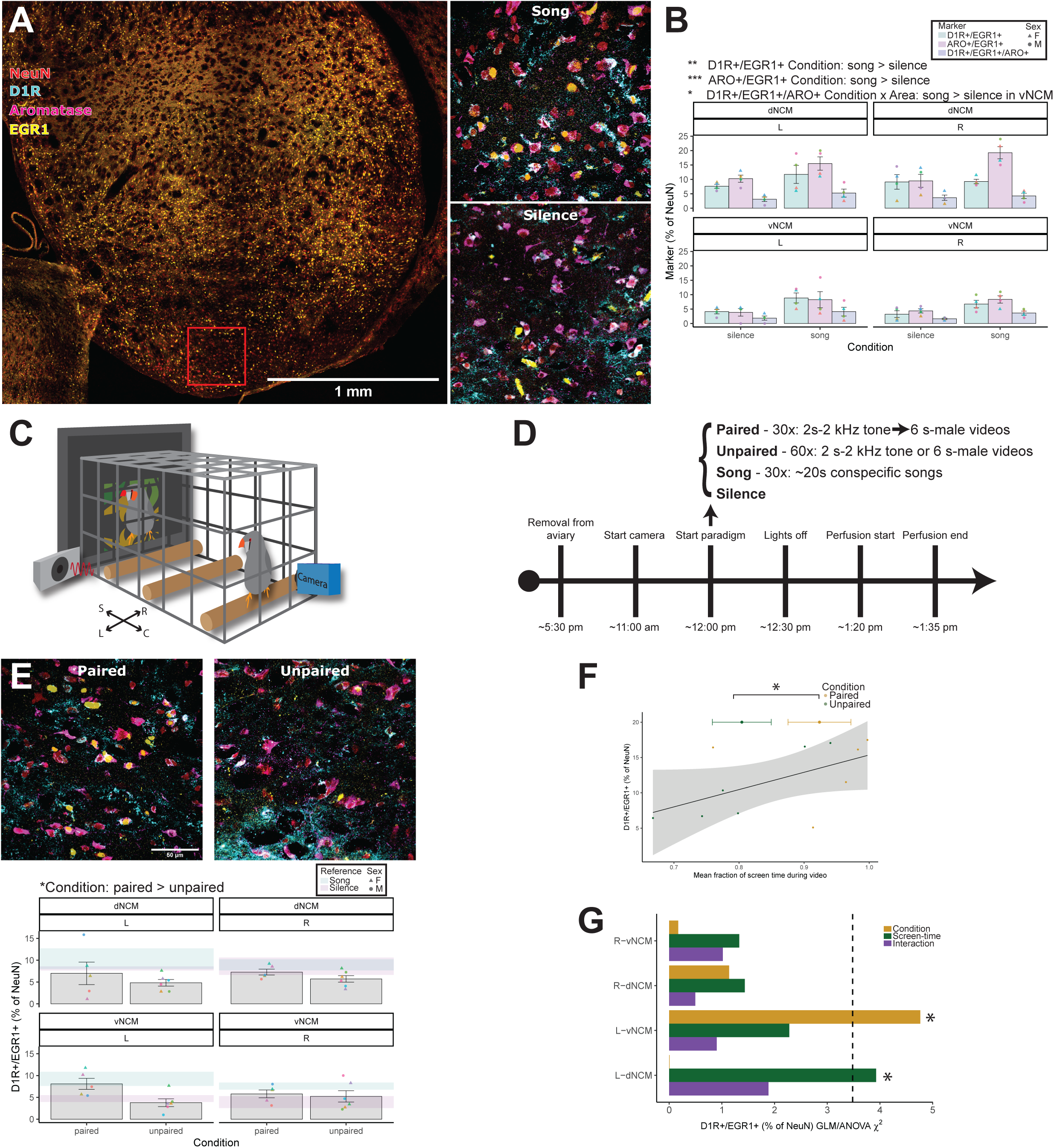
Early-growth response 1 (EGR1) protein expression in NCM neurons after song stimulation alone or auditory classical conditioning with pure tone and social reinforcement. (A-B) Song stimulation increases EGR1 expression increases in D1R+ and ARO+ across areas and hemispheres, and D1R+/ARO+ neurons in the vNCM across hemispheres. (C) Classical conditioning behavioral apparatus and (D) timeline. After behavioral manipulations, a 50 min lights-off period followed, then animals were quickly perfused and brains were processed for EGR1, D1R and aromatase (ARO) labeling (A,E). (E) D1R+/EGR1+ neurons are higher in the Paired than in the Unpaired group. Note that in left ventral NCM the Paired group mean is within the range of the Song Reference group. (F) Screen-time predicts EGR1 expression in D1R+ neurons and is higher in Paired than in Unpaired groups. (G) GLM/ANOVA χ^2^ results for the frequency of D1R+/EGR1+ neurons after classical conditioning paradigm within each hemisphere*area combination, using Condition and Scree-time as predictors. Significant effects were found for Condition and not Screen-time in the L-vNCM, and of Screen-time and not Condition in the L-dNCM (Paired > Unpaired in both). *p<0.05, **p<0.01, ***p<0.001; L: left; R: right; NCM: caudomedial nidopallium; dNCM: dorsal NCM, vNCM: ventral NCM. Marker colors represent data from individual subjects within each experiment.

### 1.2.3 Classical conditioning increases EGR1 expression specifically in D1R+ neurons

We were interested in whether our novel classical conditioning paradigm would induce expression of EGR1 in NCM neurons, so we compared expression levels between the Paired group (sound followed by video) and the Unpaired group (sound and video presented at random; Fig. 2C-D). We found that overall EGR1 neuronal induction (i.e., total EGR1 as a % of NeuN) was not affected by Condition (Supplementary Fig. 3B; Area (dNCM>vNCM): χ^2^(1)=24.726, p<0.001; Condition: χ^2^(1)=2.154, p=0.142; p>0.187 in all other main effects). However, the frequency of double-labeled D1R+/EGR1+ neurons was significantly increased by Condition (Fig. 2E; Condition: χ^2^(1)=6.370, p=0.012; p>0.150 in all other main effects). This effect was not observed with ARO+/EGR1+ neurons (Supplementary Fig. 3C; Area (dNCM>vNCM): χ^2^(1)=22.430, p<0.001; Condition: χ^2^(1)=1.027, p=0.311; p>0.290 in all other main effects) or with D1R+/ARO+/EGR1+ neurons (Supplementary Fig. 3D; Condition: χ^2^(1)=1.593, p=0.207; p>0.232 in all other main effects).

We note that we observed a significant interaction between hemisphere and condition on our NeuN counts (Supplementary Fig. 3A; Area (dNCM>vNCM): χ^2^(1)=9.910, p=0.002; Hemisphere*Condition: χ^2^(1)=6.376, p=0.012; p>0.068 in all other main effects. Tukey’s post-hoc test Paired vs Unpaired: L hemisphere: t(33)=–2.217, p=0.034, R hemisphere: t(33)=0.226, p=0.822). We ran a GLM/ANOVA including NeuN counts as a covariate to examine its impact on D1R+/EGR1+ frequency. We found that NeuN numbers did not account for the differences we observed above, i.e. that classical conditioning increased the frequency of D1R+/EGR1+ expression in NCM neurons (NeuN: χ^2^(1)=0.998, p=0.318; Condition: χ^2^(1)=6.748, p=0.009. p>0.229 in all other main effects). We are unable to explain this interaction between Condition and Hemisphere in NeuN expression in our sample. Still, this result has little bearing on our main conclusion that classical conditioning increases EGR1 expression in D1R+ neurons.

Next, we wanted to explore whether the increase in EGR1 expression in D1R+ neurons in our Paired group could be explained by any behavioral parameter. Specifically, we were concerned that differences in “reward” exposure (i.e. more time looking at the screen) could underlie the difference we observed between treatment groups. Indeed, we individually tested all behaviors we scored (Supplementary Fig. 4) and found that average screen-time (beak directions towards left, right or screen; see Methods) was significantly higher in the Paired group (GLM/ANOVA: χ^2^(1)=4.820, p=0.028; Fig. 2F; Supplementary Fig. 4F). Similarly, state behaviors that reflected distractibility towards the video (eyes closed, eating, drinking, continuously flying, beak direction towards camera) were higher in the Unpaired group (GLM/ANOVA: χ^2^(1)=10.08, p=0.001; Supplementary Fig. 4F).

To formally test whether screen-time predicted EGR1 expression in D1R+ neurons, we performed a GLM including screen-time as a covariate. Indeed, screen-time was a predictor of D1R+/EGR1+ expression, but there was a significant interaction between all factors (Fig. 2F; Screen-time: χ^2^(1)=9.816, p=0.003; Hemisphere*Area*Condition*Screen-time: χ^2^(1)=4.137, p=0.042; all other p>0.068). Distractibility did not predict D1R+/EGR1+ (Distractibility: χ^2^(1)=2.441, p=0.118; all other p>0.153).

Because we observed a 4-way interaction with hemisphere*area*condition*screen-time, we retested these data using GLMs for each hemisphere*area combination. We found that Condition (Paired vs Unpaired) and not screen-time was a significant predictor of D1R+/EGR1+ neurons in the L-vNCM (Condition: χ^2^(1)=4.765, p=0.029; Screen-time: χ^2^(1)=2.280, p=0.131; Condition*Screen-time: χ^2^(1)=0.903, p=0.342), indicating that associating sounds with reinforcement drove EGR1 in D1R-expressing neurons. Conversely, in the L-dNCM, screen-time (but not Condition) was a significant predictor of EGR1 in D1R+ neurons (Condition: χ^2^(1)=0.008, p=0.929; Screen-time: χ^2^(1)=3.926, p=0.047; Condition*Screen-time: χ^2^(1)=1.884, p=0.170). In the right hemisphere neither factor nor interactions were statistically significant (all p>0.231). ANOVA results are illustrated in Fig. 2G. In summary, screen-time was a strong predictor of EGR1 expression in D1R+ neurons, but this effect was particular to the L-dNCM. Importantly, classical conditioning increased EGR1 expression in D1R+ neurons in the L-vNCM which suggests that association during classical conditioning is hemisphere- and subregion-specific.

### 1.2.4 D1R+ neurons are predominantly GABA+

Next, we wanted to characterize the neurotransmitter profile of D1R+ cells in NCM. We found that the majority (58.7 and 64.2%) of D1R+ neurons are GABA+, and these colabeled neurons represent 21.9 and 26.4% of the neuronal population in dNCM and vNCM, respectively. The reciprocal was also true, such that 54.6% (dNCM) and 56.5% (vNCM) of GABA+ neurons contained D1R (Fig. 3A). A chi-square test analyzing these proportions showed that D1R+/GABA+ co-expression was significantly more frequent than expected (χ^2^(1)=319.660, p<0.001; Pearson’s residual=9.775, p<0.001). Similarly, we observed that the majority (∼55%) of parvalbumin+ (PV+) neurons also express D1R, combining vNCM and dNCM (Fig. 3B), and that this neurochemical phenotype represented ∼4% of the entire NCM neuronal population (Fig. 3A). The D1R+/PV+ cell population proportion is also more frequent than expected (χ^2^(1)=22.288, p<0.001; Pearson’s residual=3.470, p<0.001). We report D1R/GABA/PV anatomical distributions in Supplementary Fig. 5. These results show that D1R+ neurons are predominantly GABA+ and represent a significant subpopulation in NCM. Of note, our data show that the majority of PV+ neurons express D1Rs, which suggests this subpopulation is of particular interest for dopamine modulation of auditory processing.

**Figure 3.**
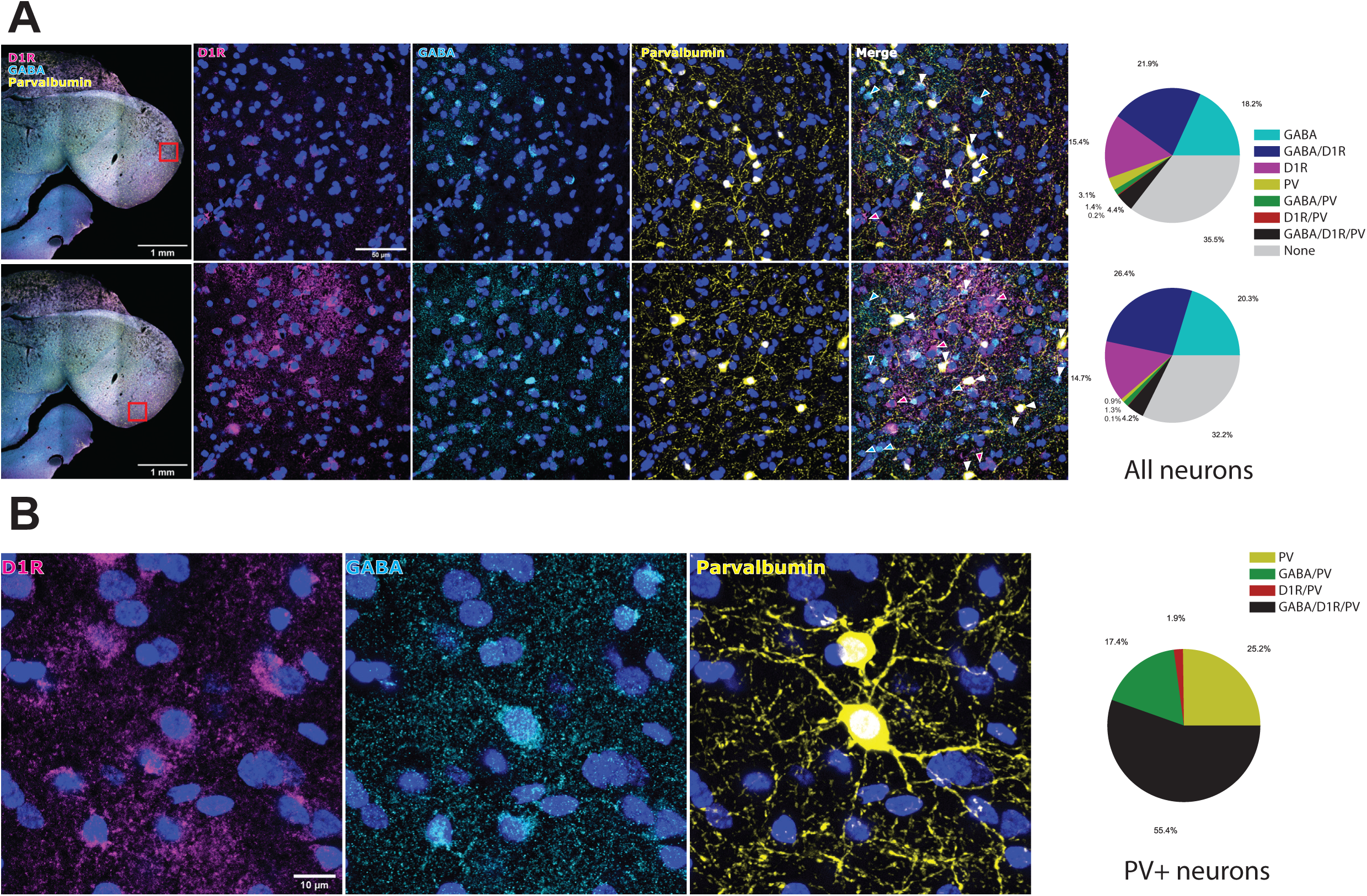
Dopamine D1 receptor (D1R)-containing neurons are predominantly GABA+. (A) Triple immunofluorescence stain for D1R, GABA and parvalbumin (PV). Magenta, cyan and yellow arrows indicate single-labeled D1R, GABA and PV+ neurons, respectively. White arrows indicate triple-labeling. (B) The majority of PV+ neurons also express GABA and D1Rs.

### 1.2.5 D1R activation reduces the amplitude of GABAergic currents in NCM *in vitro*

The anatomical findings above led to the hypothesis that activating D1Rs would modulate GABAergic neurotransmission. Thus, we recorded spontaneous postsynaptic currents from neurons in NCM *in vitro* (Fig. 4A). Inhibitory currents were isolated with the AMPA receptor antagonist DNQX (sIPSCs; Fig. 4B) and were confirmed to be GABAergic by GABA receptor antagonist bicuculline application at the end of recordings (Supplementary Fig. 6A). In separate sets of cells, we either applied 10 µM SKF or nothing (rundown) to the bath.

**Figure 4.**
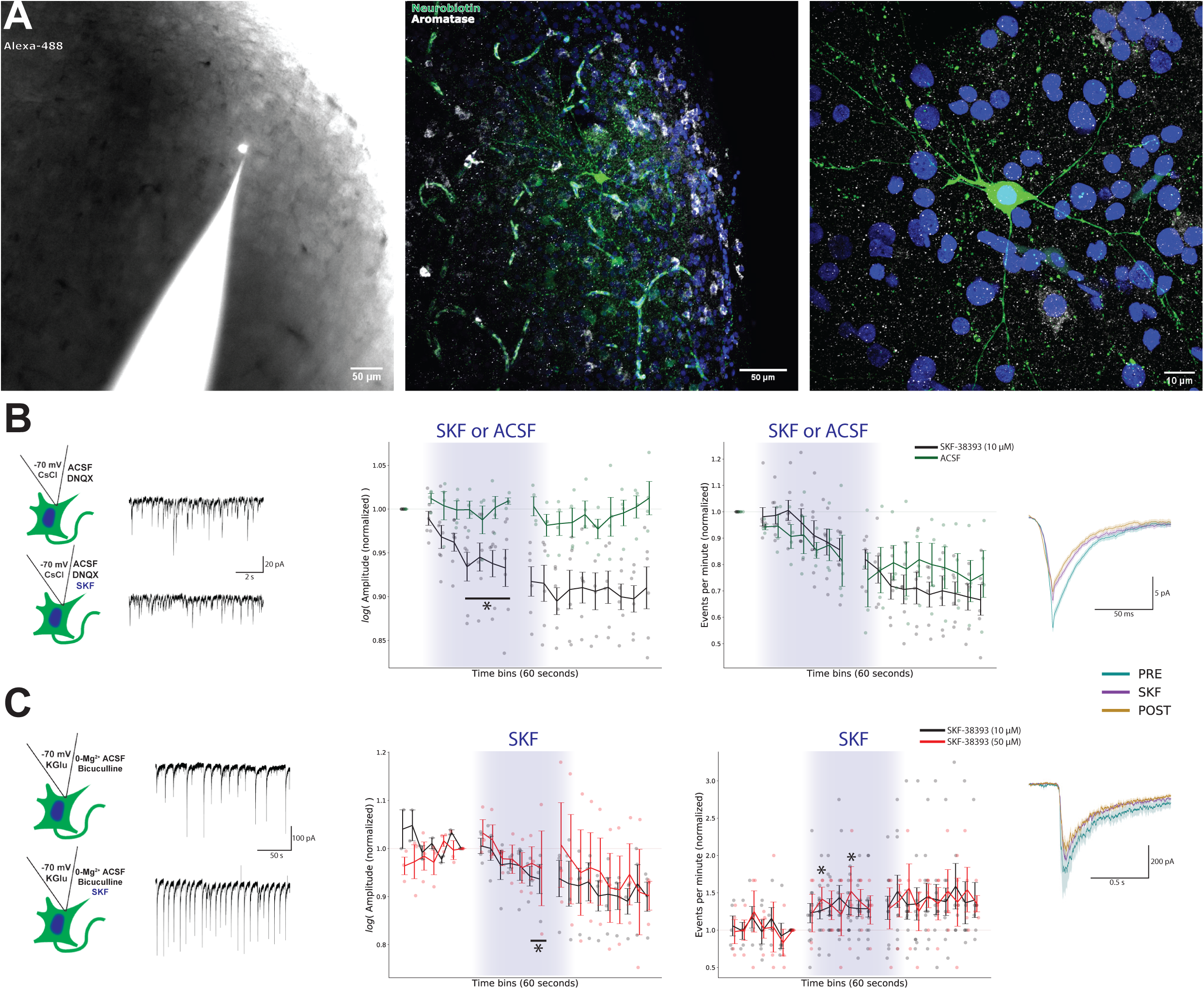
Dopamine D1 receptor (D1R) activation with SKF-38393 reduces spontaneous GABAergic (sIPSC) and glutamatergic (sEPSC) current amplitudes but increases sEPSC frequency. (A) Representative neuron in NCM visualized during (left; Alexa-488) and after recording (Neurobiotin-Streptavidin-488 stain; aromatase stain to confirm NCM location). (B) Representative sIPSCs recording from the same NCM neuron, before and during 10 µM SKF-38393 infusion (left panel). Amplitude of sIPSCs (middle panel) was significantly different from rundowns (aCSF) and reduced from baseline by SKF (4-7 min). Frequency (right panel) was also reduced but did not differ from rundown experiments (5-7 min). Waveforms show representative sIPSC (mean±SEM during 1 min) before (PRE), on the last minute of (SKF) and on the last minute after (POST) drug infusion. (C) Representative AMPA/NMDA/Kainate sEPSCs recording from the same NCM neuron, before and during 10 µM SKF-38393 infusion (left panel). Amplitude of sEPSCs (middle panel) was significantly reduced by 10 and 50 µM (no dose difference) SKF (6-7 min). In addition, frequency (right panel) was increased by SKF (2, 5 min). In these experiments, rundowns were performed before treatment and did not significantly change. Waveforms show representative sEPSC (mean±SEM during 1 min) before (PRE), on the last minute of (SKF) and on the last minute after (POST) drug infusion. *p<0.05.

For amplitude measurements, GLM/ANOVA analyses comparing treatments (SKF or rundown) detected a significant interaction between Time and Treatment (Fig. 4B; Time: χ^2^(7)=27.781, p<0.001; Treatment: χ^2^(1)=0.085, p=0.770; Time*Treatment: χ^2^(7)=15.592, p=0.029). Dunnett’s post-hoc tests revealed a significant reduction in sIPSC amplitude only in SKF treated neurons (vs before-treatment: p<0.05 between minutes 4-7 of SKF). For frequency measurements, GLM/ANOVA analyses did not detect significant differences between SKF-38393 and rundown, but detected a significant reduction of sIPSC frequency over time (Fig. 4B; Time: χ^2^(7)=27.454, p<0.001; Treatment: χ^2^(1)=0.096, p=0.757; Time*Treatment: χ^2^(7)=1.735, p=0.973; Dunnett’s post-hoc test vs before-treatment: p<0.05 between minutes 5-7). These results show that SKF-38393 treatment significantly reduced the amplitude of GABAergic currents. The reduction in frequency observed during SKF-38393 treatments did not differ from rundowns indicating that, regardless of treatment, the number of detected sIPSCs decayed over time.

### 1.2.6 D1R activation reduces the amplitude but increases the frequency of glutamatergic currents in NCM *in vitro*

Our anatomical findings suggest that ∼38% of D1R-positive neurons are non-GABAergic (i.e. putatively glutamatergic; see Fig. 3A), which led to the hypothesis that activating D1Rs would also directly modulate glutamatergic currents in NCM. Therefore, we recorded excitatory NMDA/AMPA/Kainate currents (sEPSCs) isolated in 0-Mg^2+^ bath containing the GABA_a_- receptor antagonist bicuculline (Fig. 4C). These currents were confirmed to be NMDA dependent at the end of the recordings (Supplementary Fig. 6B). For these experiments we used two doses (10 and 50 µM) of SKF-38393 in different sets of cells. In these experiments we performed rundown experiments before SKF-38393 treatment in a subset of cells (n=4 for amplitude; n=8 for frequency; see methods). Both amplitude and frequency of sEPSCs were stable during 7 minutes before treatment (Fig. 4C; GLM/ANOVA: Amplitude: χ^2^(6)=5.056, p=0.537; Frequency: χ^2^(6)=8.098, p=0.231).

For amplitude measurements following SKF-38393 treatment, we performed GLM/ANOVA on the effect of different doses of SKF-38393 over time. These analyses showed a reduction in the amplitude of sEPSCs over time due to treatment, and no difference between the two SKF doses (Fig. 4C; Time: χ^2^(7)=41.458, p<0.001; Treatment: χ^2^(1)=0.100, p=0.752; Time*Treatment: χ^2^(7)=1.963, p=0.962; Dunnett’s post-hoc test vs before-drug: p<0.05 on minutes 6-7 of SKF). For frequency measurements, GLM/ANOVA analyses showed an increase in sEPSC frequency over time, and again no difference between the two SKF doses (Fig. 4C; Time: χ^2^(7)=21.802, p=0.003; Treatment: χ^2^(1)=2.480, p=0.115; Time*Treatment: χ^2^(7)=4.000, p=0.780; Dunnett’s post-hoc test vs before-drug: p<0.05 on minutes 2 and 5 of SKF).

In summary, D1R activation *in vitro* reduced the amplitude of both GABA and glutamatergic spontaneous currents, while also increasing the frequency of the latter. These findings establish a role for dopamine modulation of network excitation and inhibition in NCM, and predict that D1R activation *in vivo* causes differential effects depending on cell type (i.e., downregulate vs. upregulate GABAergic and glutamatergic neuron firing, respectively).

### 1.2.7 Cell type separation based on waveform measurements for *in vivo* recordings

We isolated 107 single-units from 9 adult birds in awake head-fixed recordings using acute recording 32-channel microdrives coupled to retrodialysis probes (RetroDrives; Supplementary Fig. 7; see methods). We measured peak-to-peak duration and ratio of each unit and analyzed the data using an unsupervised hierarchical clustering algorithm (see methods; Fig. 5A). The gap-statistic results show that the variance in clustering was best explained by 4 clusters. Cell types differed significantly in waveform peak-to-peak duration (GLM/ANOVA: χ^2^(3)=299.970, p<0.001; Tukey’s post-hoc test: all p<0.01) and ratio (GLM/ANOVA: χ^2^(3)=294.880, p<0.001; Tukey’s post-hoc test: all p<0.001 except BS1–BS2 where p=0.972). In summary, waveform peak-to-peak duration followed the pattern NS1<NS2<BS1<BS2 and waveform peak-to-peak ratio followed BS2<BS1=NS1<NS2.

**Figure 5.**
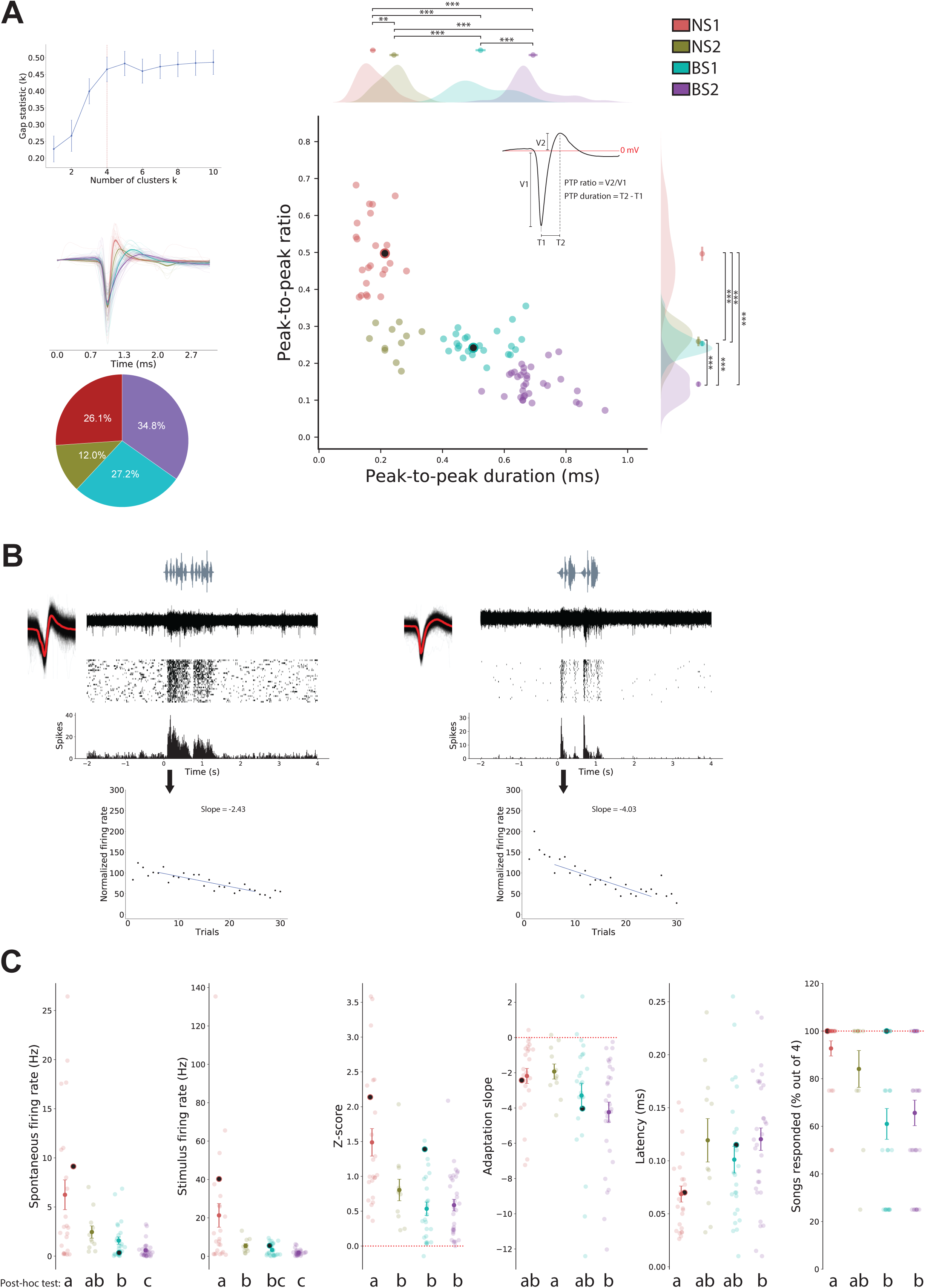
Cell type separation based on waveform measurements in *in vivo* recordings. (A) Unsupervised hierarchical clustering results (gap statistic) using peak-to-peak ratio vs duration (top left); separation into 4 clusters best explained the variance in the data. Traces illustrate all waveforms in our sample by cell type (middle left). Shaded lines are the average waveforms of each unit in our sample; opaque lines and shading are the grand average±SEM of those. The pie chart represents the proportions of units within each cluster in our sample (bottom left). The correlation plot shows the 4 distinct clusters and the kernel-density estimations of the distributions along single-axis with mean±SEM on top. (B) Representative PSTHs and adaptation slopes of NS1 (left) and BS1 (right) cell types. Representative examples in B are highlighted in all graphs according to cluster type. (C) Physiology parameters of the different cluster types. From left to right: spontaneous and stimulus firing rate, z-score, adaptation slope, latency to respond to song and percentage of songs responded to. Post-hoc test results are shown at the bottom. Significant differences (p<0.05) are assigned different letters. *p<0.05, **p<0.01, ***p<0.001. Highlighted datapoints in A,C depict the two representative cells in B.

The classification commonly used in the literature of narrow- and broad-spiking neurons in songbird pallium uses only peak-to-peak duration and a division boundary of ∼0.4 ms (Aurore et al., 2019; Schneider and Woolley, 2013; Vahaba et al., 2017; Yanagihara and Yazaki-Sugiyama, 2016). Using both peak-to-peak duration and ratio, we provide evidence of 2 further subdivisions; therefore, we named our clusters to extend the previous classification: NS1 and NS2 – narrow-spiking; and BS1 and BS2 – broad-spiking.

Following clustering, non-auditory-responsive cells were excluded from the analyses (see methods) and 92 units were further analyzed. This sample consisted of 21 (L) and 23 (R) units from females and 34 (L) and 14 (R) units from males. In this sample, BS2 were more numerous (34.8%) followed by BS1 (27.2%), NS1 (26.1%) and NS2 (12%; Fig 5A). Representative PSTHs and adaptation slopes of a NS1 and a BS1 are shown in Fig. 5B.

The baseline physiological profiles assessed during aCSF infusion showed clear segregation by cell type classification (Fig. 5C). GLM/ANOVA analyses showed that cell types differed in spontaneous firing rates (χ^2^(3)=50.554, p<0.001; Tukey’s post-hoc test: p<0.05 in NS1–BS1, NS1–BS2, NS2–BS2 and BS1–BS2), stimulus firing rates (χ^2^(3)=86.754, p<0.001; Tukey’s post-hoc test: p<0.05 in NS1–NS2, NS1–BS1, NS1–BS2 and NS2–BS2), z-scores (χ^2^(3)=36.169, p<0.001; Tukey’s post-hoc test: p<0.05 in NS1–NS2, NS1–BS1 and NS1–BS2), adaptation slopes (χ^2^(3)=10.465, p=0.015; Tukey’s post-hoc test: p<0.05 in NS2–BS2), latencies to respond (χ^2^(3)=12.166, p=0.006; Tukey’s post-hoc test: p<0.05 in NS1–BS2) and in the % of songs responded to (χ^2^(3)=22.785, p<0.001; Tukey’s post-hoc test: p<0.05 in NS1–BS1 and NS1–BS2). Therefore, the 4 cell types clustered by waveform shape in our recordings also differed in physiological profile.

Broadly speaking, NS1 cells had highly symmetrical and narrow action potentials, high firing rates and z-scores, as well as fast response latencies and lower selectivity, which all parallel features of mammalian cortical high-firing PV+ inhibitory interneurons. Compared to NS1, NS2 cells had less narrow and symmetrical waveforms with lower firing rates, which resemble properties of mammalian cortical low-firing somatostatin+ or VIP+ inhibitory interneurons (Tremblay et al., 2016). BS1 and BS2 had broader and less symmetrical waveforms and lower firing rates than NS1/NS2. Additionally, they showed longer latencies to respond to stimuli and higher selectivity, all features that resemble those of mammalian cortical excitatory projection neurons (Atencio and Schreiner, 2008; Hromádka et al., 2008; Tsunada et al., 2012; Wu et al., 2008).

### 1.2.8 D1R activation reduces spontaneous and stimulus firing of NS1, but increases spontaneous firing of BS1 neurons *in vivo*

Our *in vitro* experiments generated clear predictions about differential D1R activation on narrow- vs. broad-spiking neurons *in vivo* (see above). We therefore analyzed how SKF-38393 (SKF; 0.2 mM) affected single-unit responses to sound playback (timeline in Fig. 6). Representative PSTHs for NS1 and BS1 cells are shown in Fig. 7A. We observed effects of D1R activation that were selective for cell waveform phenotypes in NCM.

**Figure 6.**
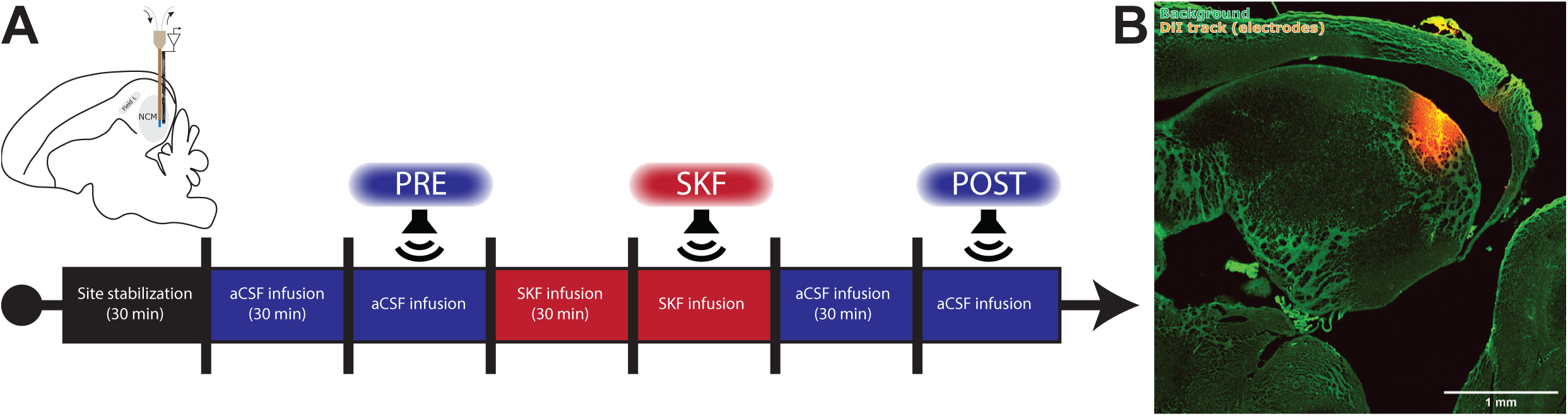
(A) Timeline for drug infusions and playbacks starting after auditory site localization within NCM. (B) Electrode track confirmation. Before insertion in the brain, electrodes were dipped in DiI-594 (6.25% in 200-proof ethanol) for posthumous site confirmation.

**Figure 7.**
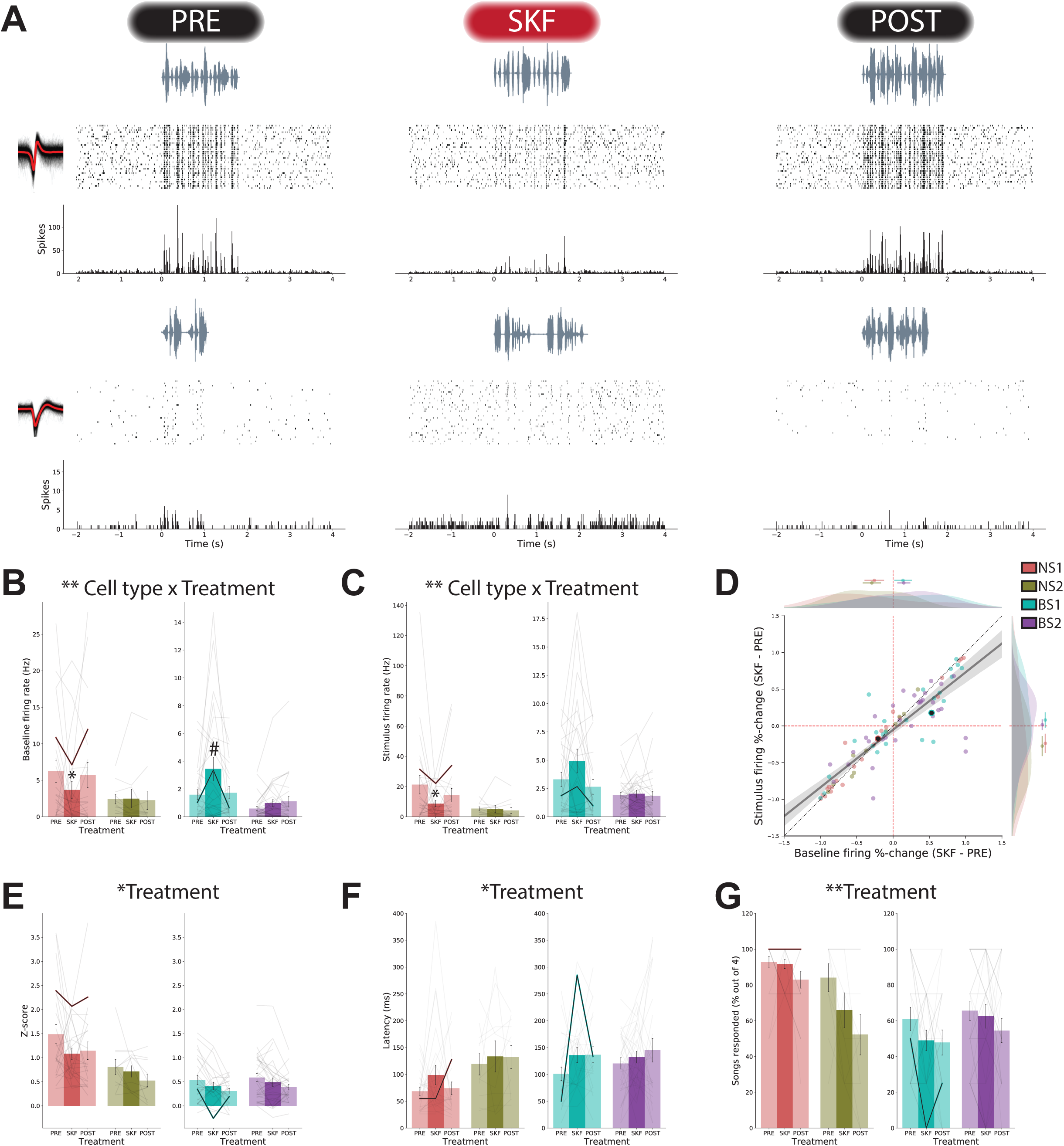
Effects of D1R activation *in vivo* on firing characteristics of NCM neurons. (A) Representative PSTHs of an NS1 (top) and a BS1 (bottom) cell in response to SKF. Note the change in stimulus and spontaneous firing in NS1 and BS1, respectively, due to SKF-38393 (0.2 mM). Representative cells are highlighted in all plots according to their cell type. (B) Spontaneous firing rate. D1R agonist reduced firing of NS1, but increased firing of BS1 cells (trend in full GLM model, but significant in restricted model). (C) Stimulus firing rate. D1R agonist reduced firing of NS1 cells. (D) Correlation between spontaneous and stimulus %- change during SKF infusion accompanied by kernel-density estimations of the distributions along single-axis with mean±SEM on top. Regression line across all data points is shown (solid line). Identity line is shown as a dotted line. The regression slope 95% confidence interval (shaded area) is less than and does not encompass 1, indicating that spontaneous firing changes are higher than stimulus firing changes. (E) D1R agonist reduced Z-score, (F) increased latency and (G) decreased responsiveness of NCM cells. Relevant statistical effects are highlighted on top of each plot. Post-hoc Tukey test results are displayed on the plots when interaction was significant. ^#^p<0.1; *p<0.05; **p<0.01.

For spontaneous firing rate, GLM/ANOVA analyses comparing Treatment and Cell-Type showed that SKF reduced the firing of NS1 and tended to increase the firing of BS1 (Fig. 7B; Treatment: χ^2^(1)=0.011, p=0.916; Cell-Type: χ^2^(3)=54.694, p<0.001; Treatment*Cell-Type: χ^2^(3)=12.866, p=0.005; Tukey’s post-hoc test: PRE–SKF: NS1: t(173)=2.702, p=0.007; NS2: t(173)=0.712, p=0.477; BS1: t(173)=–1.758, p=0.079; BS2: t(173)=–1.155, p=0.248). To assess whether a treatment effect could be detected in BS1 units with increased statistical power, we performed a single-factor GLM/ANOVA only on BS1 units, which yielded a significant increase in spontaneous firing due to SKF (χ^2^(1)=6.008, p=0.014).

For stimulus firing rate, SKF decreased firing of NS1 cells (Fig. 7C; Treatment: χ^2^(1)=0.734, p=0.392; Cell-Type: χ^2^(3)=53.731, p<0.001; Treatment*Cell-Type: χ^2^(3)=9.043, p=0.029; Tukey’s post-hoc test: PRE–SKF: NS1: t(173)=2.753, p=0.007; NS2: t(173)=0.783, p=0.435; BS1: t(173)=–1.232, p=0.220; BS2: t(173)=–0.183, p=0.855).

To explore further the change in spontaneous and stimulus firing due to SKF, we plotted a correlation between the %-change in spontaneous versus stimulus firing induced by SKF (Fig. 7D). Values above 0 in either axis indicate an increase in firing due to SKF. Note that on average BS1 and BS2 data points were situated above 0 in both axes, whereas NS1 and NS2 were below 0. Changes in spontaneous vs stimulus firing were also highly correlated (Pearson’s r=0.876, p<0.001). Interestingly, the regression line slope’s 95% confidence interval [0.693; 0.875] falls lower than and does not include the slope of the identity line (slope=1), which suggests that spontaneous firing was more affected by SKF than stimulus firing.

For stimulus response z-scores, GLM/ANOVA analyses showed that SKF decreased overall z-scores regardless of cell type (Fig. 7E; Treatment: χ^2^(3)=8.657, p=0.003; Cell-Type: χ^2^(3)=46.855, p<0.001; Treatment*Cell-Type: χ^2^(3)=4.443, p=0.217).

Finally, SKF-38393 treatment increased overall latency to respond (Fig. 7F; Treatment: χ^2^(3)=5.754, p=0.016; Cell-Type: χ^2^(3)=13.597, p=0.004; Treatment*Cell-Type: χ^2^(3)=2.248, p=0.523) and decreased the % of songs units responded to (Fig. 7G; Treatment: χ^2^(3)=8.929, p=0.003; Cell-Type: χ^2^(3)=31.935, p<0.001; Treatment*Cell-Type: χ^2^(3)=4.409, p=0.221) regardless of cell type.

Altogether, these results show that the D1R agonist SKF-38393 affects the response properties of NCM cell types differentially. Notably, as predicted by our *in vitro* experiments, D1R activation reduces the firing of putative inhibitory neurons (NS1) while increasing the firing of putative excitatory neurons (BS1). Furthermore, in all cell types D1R activation decreased z-scores, increased latency to respond and decreased the % of songs units responded to, which might represent direct or indirect consequences of the shift in inhibitory/excitatory balance induced by tonic D1R activation.

### 1.2.9 D1R activation sharply reduces neuronal stimulus-specific adaptation *in vivo*

NCM neurons show stimulus-specific adaptation when birds are presented repetitions of the same stimuli, which are thought to reflect short/medium-term memory formation (Chew et al., 1996; Lu and Vicario, 2017). Therefore, we asked whether D1R activation would result in changes in adaptation to novel stimuli. Trial-by-trial stimulus firing rates were used for deriving adaptation slopes (see methods). Eight cells (3 BS1, 5 BS2) had to be removed from the analyses, because firing rate on the first trial used for the regression (trial 6; see methods) was 0 during either PRE or SKF playbacks. Representative rasters and corresponding slopes of an NS1 cell is shown in Fig. 8A. Fig. 8B depicts the slope through the normalized firing rate of trials 6-25 averaged by cell type, for all cell types. GLM/ANOVA analyses showed that SKF-38393 infusion reduced adaptation (i.e. shallower regression slopes), regardless of cell type (Fig. 8C; Treatment: χ^2^(3)=8.278, p=0.004; Cell-Type: χ^2^(3)=20.069, p<0.001; Treatment*Cell-Type: χ^2^(3)=5.244, p=0.155). Note, however that this effect seems to be driven by all cell types but for BS2, which on average appears to remain unchanged by SKF. In fact, BS2 cells were the only group that retained non-zero slopes during SKF treatment (One-sample Wilcoxon signed-rank tests versus 0; NS1: p=0.308; NS2: p=0.549; BS1: p=0.052; BS2: p<0.001).

**Figure 8.**
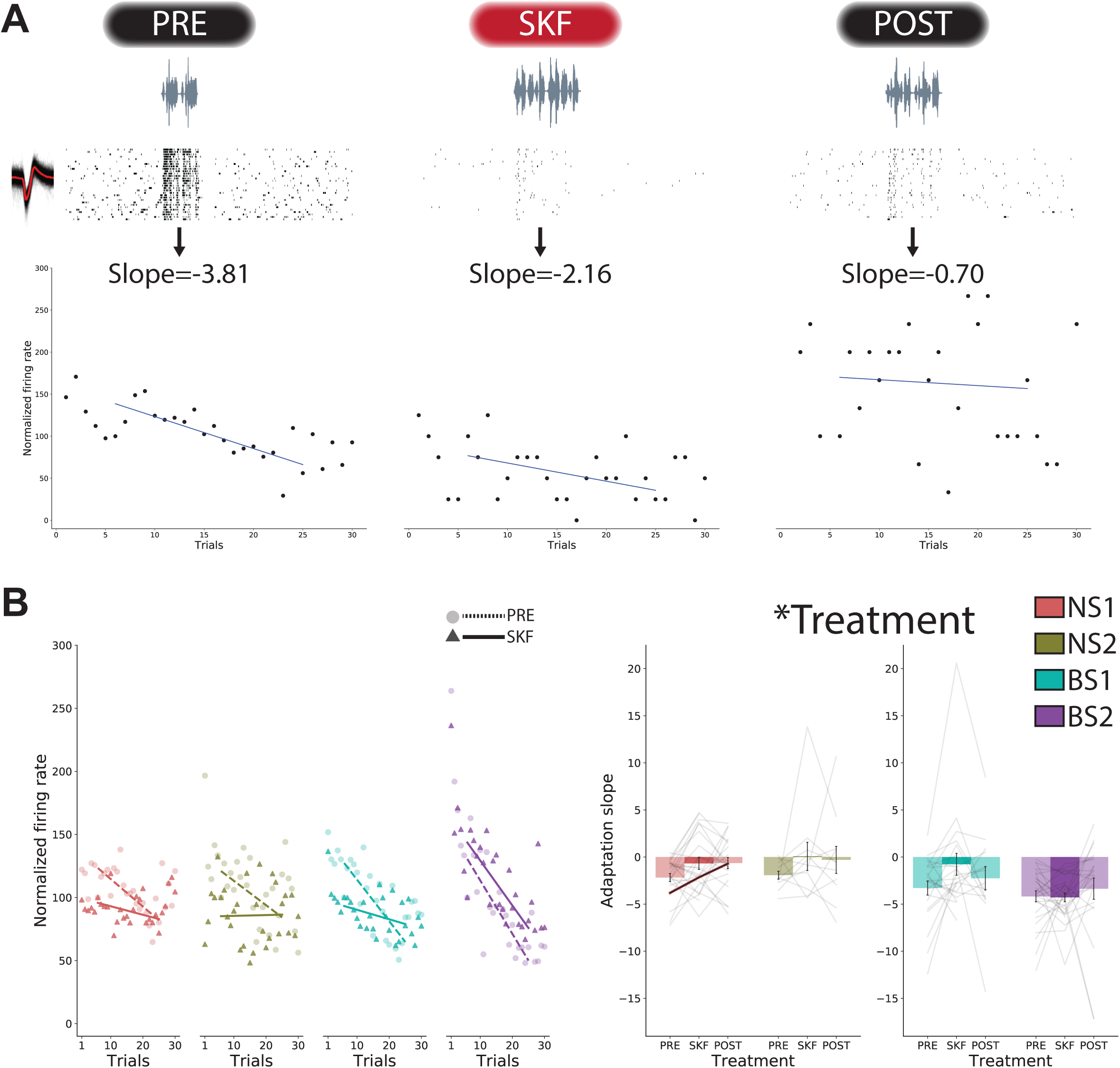
D1R activation disrupts neuronal stimulus-specific adaptation *in vivo*. (A) Representative PSTHs from a single NS1 highlighting a decrease in adaptation slope to a new stimulus due to SKF-38393 (0.2 mM) treatment. (B) The slope through the normalized firing rate of trials 6-25 averaged by cell type. (C) Adaptation slopes quantifications show a reduction in the slope due to SKF-38393. Relevant statistical effect is displayed on the top of the plot. *p<0.05.

Importantly, the %-change in slopes did not correlate with the %-change in spontaneous firing (GLM/ANOVA; Spontaneous %-change: χ^2^(1)=1.126, p=0.289; Cell-Type: χ^2^(3)=2.723, p=0.436; Spontaneous %-change*Cell-Type: χ^2^(3)=3.451, p=0.327) or stimulus firing (GLM/ANOVA; Stimulus %-change: χ^2^(1)=0.800, p=0.371; Cell-Type: χ^2^(3)=2.764, p=0.430; Stimulus %-change*Cell-Type: χ^2^(3)=2.765, p=0.429), suggesting that lower slopes are not predicted by firing rate “floor-effect” (data not shown).

Therefore, D1R activation flattens the stimulus-specific adaptation slopes of NCM neurons, consistent with the hypothesis that adaptation and memory formation in NCM are governed by the local activation of D1Rs.

## 1.3 Discussion

In this study, we show that dopamine D1 receptors (D1R) modulate learning and synaptic plasticity in the secondary association pallium (NCM) of a songbird. Specifically, we show that (1) D1R protein is prevalent in NCM neurons, especially in aromatase-, GABA-, and parvalbumin-positive neurons; (2) song stimulation increases the immediate early gene EGR1’s expression in D1R+, aromatase+ and double-labeled D1R+/aromatase+ neurons; (3) auditory classical conditioning increases EGR1 expression in D1R+ neurons; (4) activating D1Rs *in vitro* reduces the amplitude of GABAergic currents and decreases the amplitude but increases the frequency of glutamatergic currents; (5) activating D1R *in vivo* reduces firing of putative-inhibitory interneurons, while increases firing of putative-excitatory projection neurons; and (6) D1R activation *in vivo* disrupts stimulus-specific adaptation in NCM neurons, a phenomenon reflective of auditory memory formation. Together, these parallel pieces of evidence are consistent with the hypothesis that dopamine is an important modulator of complex sensory function and plasticity and a possible substrate for reinforcement learning. We discuss the significance and broader implications of our findings below.

Dopamine signaling modulates auditory association learning in primary auditory cortex (Bao et al., 2001; Reichenbach et al., 2015; Schicknick et al., 2012, 2008). In these studies, dopamine activation was paired with simple stimuli such as pure and frequency-modulated tones. However, there is limited work on the role of dopamine in modulating the processing of complex auditory signals, in high-order cortical/pallial structures. In humans, systemic dopamine-enhancing treatments enhanced auditory language learning (Breitenstein et al., 2004; Knecht et al., 2004). Here, we show in a songbird that dopamine signaling specifically in high-order sensory pallium can modulate the encoding of complex auditory signals by shifting inhibitory-excitatory balance, in favor of the hypothesis of sustained spontaneous excitation by means of disinhibition.

A distinct feature of NCM among the auditory forebrain nuclei is the dense labeling for somatic aromatase, an enzyme that mediates conversion of testosterone into estradiol (E2) (Saldanha et al., 2000). E2 production in NCM is elevated during social interactions and song playbacks (Remage-Healey et al., 2012, 2008), and E2 treatment rapidly increases neuronal response to song playbacks (Remage-Healey et al., 2010). We recently showed that blocking aromatase locally in NCM slows auditory association learning in an operant task (Macedo-Lima and Remage-Healey, 2020). This finding led to the hypothesis that dopamine interacts with E2 signaling in NCM to support association learning. This line of reasoning is also supported by reports that dopamine in the auditory cortex mediates auditory association learning in gerbils (Schicknick et al., 2012), and that dopamine innervation and release are increased by steroid hormones in songbird NCM (Matragrano et al., 2011; Rodríguez-Saltos et al., 2018). In the present study, we provide anatomical evidence to support this hypothesis, since ∼33% of aromatase+ neurons coexpress D1R protein and these co-labeled neurons represent ∼8% of all neurons in NCM (Fig. 1C). Interestingly, in striatum and preoptic area, E2 and dopamine systems cross-modulate (Balthazart et al., 2002; Becker, 1990; Lammers et al., 1999; Tozzi et al., 2015) and dopamine and E2 have been suggested to cross-activate each other’s receptors (Olesen and Auger, 2008; Tozzi et al., 2015). Our data suggest that in NCM E2 and dopamine signaling are acting in tandem to modulate learning and memory in the songbird auditory pallium. Thus, we hypothesize that this co-modulatory signaling could apply to other brain circuits that domiciliate both aromatase and dopamine receptors, such as human auditory cortex (Yague et al., 2006).

In songbirds, conspecific song-driven EGR1 expression is a hallmark of NCM in contrast to surrounding regions, and can even be used to define NCM’s anatomical boundaries (Mello and Clayton, 1994). Furthermore, this induction is sensitive to E2 (Krentzel et al., 2019; Lampen et al., 2017; Maney et al., 2006) and conspecific song playback increases local E2 production in NCM (Remage-Healey et al., 2012, 2008). Here, we show that song stimulation increases EGR1 expression in D1R+, ARO+ and double-labeled D1R+/ARO+ neurons, i.e. in neurons that directly participate in dopamine and E2 signaling. The functional significance of this phenomenon has not been explored in vertebrates. Specifically, is EGR1 expression directly linked to changes in ARO activity? Is EGR1 expression in D1R+ neurons dependent on dopamine release? Future work should address these questions.

EGR1 has been interpreted as a surrogate for neuronal activation in NCM (Lampen et al., 2017; Nordmann et al., 2020; Scully et al., 2017). However, engagement in a non-auditory food-aversion learning task has been shown to induce NCM EGR1 (Tokarev et al., 2011), which suggests EGR1 induction in NCM is associated with multisensory learning. Similarly, here, we show that EGR1 is enhanced specifically in D1R+ neurons after an audio-visual classical conditioning paradigm. Therefore, our findings support the view that EGR1 induction reflects neuronal plasticity, rather than simply cellular activation per se (Duclot and Kabbaj, 2017).

We acknowledge the caveat that, in our task, auditory conditioning (video always preceded by sound) prompted the Paired group to attend more to the video (i.e. more actively looking at the screen or ‘screen-time’), which was detected via our full GLM model. Still, the same analyses revealed that this effect was dependent on the area and the hemisphere analyzed. Follow-up analyses identified an effect of conditioning independent of screen-time specifically in the left ventral NCM, and remarkably, that EGR1 induction was higher specifically in D1R+ neurons (and not in all, ARO+ or D1R+/ARO+ neurons). Future studies should investigate whether dopamine signaling, and midbrain dopaminergic nuclei are required to elicit these responses.

Inhibition fundamentally controls neuronal responses to sounds (Razak and Fuzessery, 2009; Wang et al., 2002) and is required for associative learning in mammal auditory cortex (Letzkus et al., 2011). Dopamine receptors are prevalent in mammalian cortical inhibitory interneurons, especially in parvalbumin-expressing neurons (Le Moine and Gaspar, 1998). Here, we show in the songbird NCM that D1R and inhibitory markers are frequently coexpressed in neurons, representing more than half of the total GABA+ and D1R+ population, and about 24% of the total NCM population.

Parvalbumin-positive (PV+) cortical interneurons are generally characterized by their fast action potentials and high sustained firing frequency and play a central role in regulating microcircuit activity states and learning processes in hippocampus and cortex (Cardin, 2018; Donato et al., 2013). In the songbird song control circuit, PV+ neurons are recruited during singing, and PV+ neuron numbers correlate with critical period closure for song learning (Balmer et al., 2009; Yildiz and Woolley, 2017). In the NCM, PV and calbindin seem to be expressed in different neuron subpopulations, such that PV+ but not calbindin+ neurons can express aromatase (Ikeda et al., 2017). Here, we show that PV+ neurons represent ∼8% of all NCM neurons, and ∼11% of GABA+ neurons. Furthermore, 55% of the parvalbumin-expressing neurons in NCM also express D1R. We hypothesize that many of the effects we observed *in vivo* and *in vitro* can be attributed to this subpopulation. This hypothesis is also supported by the waveform phenotype of the NCM neurons we observe are sensitive to SKF-38393 (see below).

D1Rs are generally assumed to increase circuit excitability through Gα_s_-protein coupling (Beaulieu and Gainetdinov, 2011). In our *in vitro* experiments, D1R agonist SKF-38393 caused a seemingly counterintuitive reduction in the amplitude of GABAergic and glutamatergic currents. However, D1R-mediated depression is well documented in the mammalian nucleus accumbens for both GABAergic and glutamatergic (especially NMDA-mediated) synapses and is attributed to presynaptic plasticity (Pennartz et al., 1992; Zhang et al., 2014). In contrast, in mammalian cortex, excitability-reducing effects are typically attributed to postsynaptic D2R-mediated effects, while D1Rs mediate excitability increases (Darvish-Ghane et al., 2016; Gonzalez-Islas and Hablitz, 2003). In our *in vitro* proposed model (Fig. 9A), for the reduction of GABA release we suggest two scenarios: 1) D1Rs are predominantly mediating an increase in GABAergic tone by neurons upstream to those providing input to the recorded neuron, therefore reducing GABA release downstream and 2) D1Rs are acting directly on neurons providing input to the recorded neuron causing a direct reduction in GABA release. For the reduction of glutamatergic current amplitude, we suggest a presynaptic mechanism, perhaps explained by a depletion in presynaptic glutamate stores, resulting from the increased firing (Staley et al., 1998), regulation of glutamate production (Sherman and Mott, 1985), or conversion of glutamate into GABA by glutamine decarboxylase (Neary et al., 1972) by D1Rs. Alternatively, SKF-38393 could be acting directly on the recorded neuron (postsynaptically) in NCM to result in amplitude reductions. Further experiments with miniature events or with synaptic stimulation could help clarify these questions.

**Figure 9.**
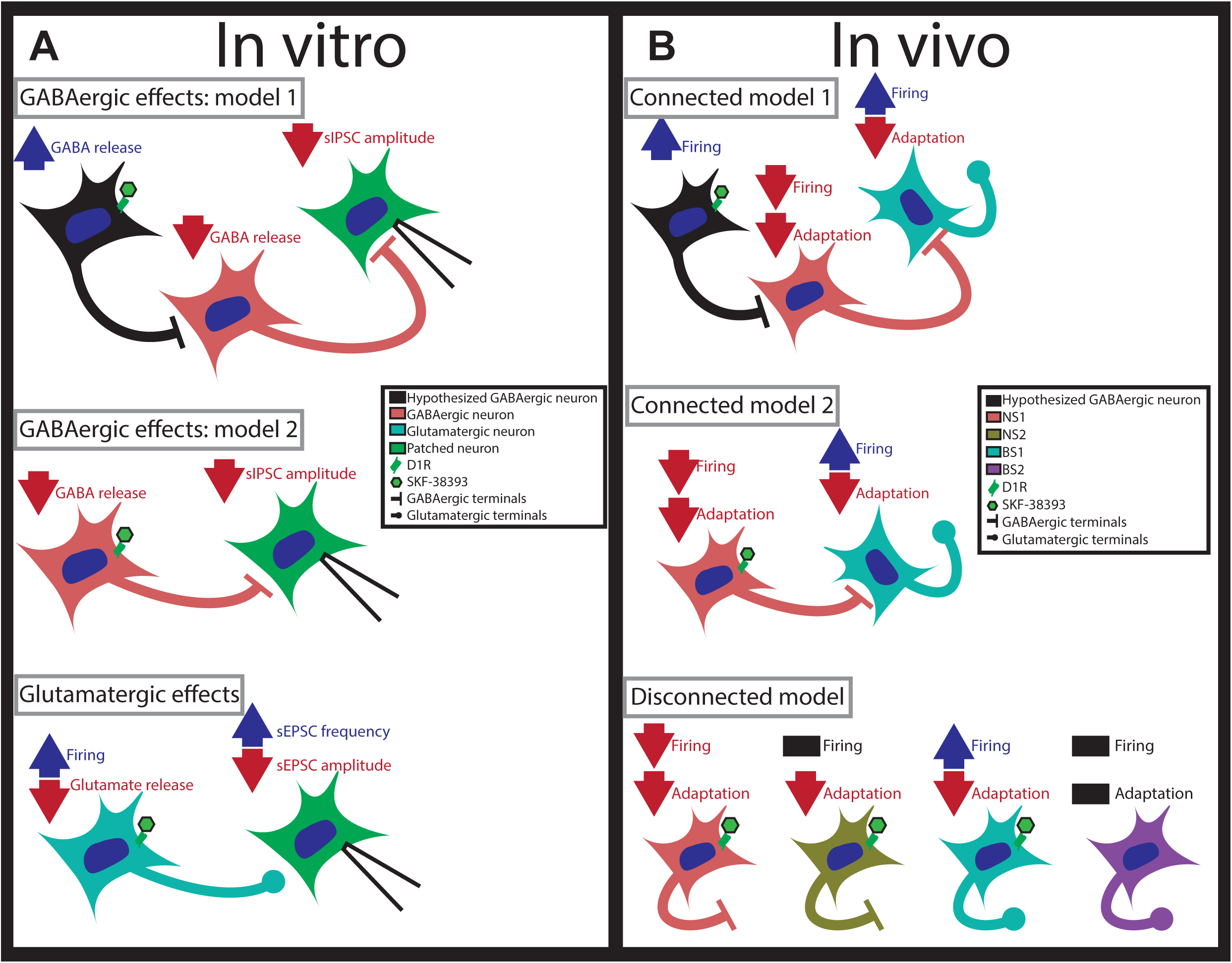
Network connectivity models hypothesized from current findings. SKF-38393 effects are shown by the large arrows. (A) Models for *in vitro* results with sIPSCs (GABAergic) and sEPSCs (glutamatergic). GABAergic model 1 hypothesizes that D1R agonist binds preferentially to a GABAergic neuron upstream from the ones providing input to the recorded neuron. Binding of D1R agonist increases its GABA tone resulting in a reduction of GABA release immediately upstream of the recorded neuron. GABAergic model 2 suggests D1R effects directly cause a reduction in GABA release by the upstream neuron. Glutamatergic effects can be explained by a direct effect of D1R agonist on a glutamatergic neuron directly upstream from the recorded neuron, causing a reduction (perhaps a depletion) of glutamate release but an increase in firing rate. (B) Models for *in vivo* results. Connected models only depict cell types with changes in firing rate. Model one hypothesizes that D1R agonist binds to and excites a GABAergic neuron upstream of NS1 cells, which results in a reduction of NS1 firing and consequent disinhibition of BS1 cells. Model 2 offers a direct effect of D1R agonist on NS1 cells, resulting in a reduction in their firing, which results in disinhibition of BS1 cells. In the disconnected model, the effects observed here are due to D1R’s binding to all cell types individually (except BS2, in which no effects were observed). sI/EPSC: spontaneous inhibitory/excitatory postsynaptic current.

We are at the beginning of our understanding of how non-layered cortical-like microcircuits operate, including their component cell types. Most of the electrophysiology studies reporting *in vivo* waveform shape segregation in NCM have relied on a non-statistical split of peak-to-peak ratio at ∼0.4 ms to divide recorded waveforms into the categories narrow- (NS) and broad-spiking (BS) units (Aurore et al., 2019; Schneider and Woolley, 2013; Vahaba et al., 2017; Yanagihara and Yazaki-Sugiyama, 2016). One study using intracellular sharp-electrode recordings has reported 4 subtypes based on visual inspection of waveform shape and firing rates (Bottjer et al., 2019). Our findings with unsupervised hierarchical clustering using solely waveform shapes generally corroborates the latter finding, such that their two NS types displayed high and low firing rates, similar to our NS1 and NS2 respectively. Our two BS subtypes, however, are contained within the range of one of their BS subtype measurements. Their other BS subtype (termed double-trough) was not representative in our sample, possibly a limitation of extracellular recordings.

The firing characteristics of the cell types identified here led us to formulate hypotheses based on characteristics of mammalian cortical neurons. NS1 neurons are highly reminiscent of mammalian fast-spiking PV+ interneurons, exhibiting short and symmetrical action potentials, high firing, short latency to respond and low stimulus selectivity (Atallah et al., 2012; Tremblay et al., 2016). Because of the narrower waveform, lower firing rate, and longer onset latencies, we suggest that NS2 resemble late-spiking interneurons, such as somatostatin+ or VIP+ interneurons (Tremblay et al., 2016). BS1 and BS2 present distinct waveforms, BS1 has a higher spontaneous firing rate than BS2 and BS2 is not remarkably sensitive to SKF-38393. We found ∼33% of NCM neurons do not express either GABA or D1Rs (Fig. 3A). Therefore, it is possible that BS2 cells are part of a circuit in which D1R-signaling does not participate to produce strong effects on variables analyzed in this study.

Alternatively, some synaptic plasticity effects can require simultaneous D1- and D2- family receptor activation (Calabresi et al., 1992; Ichihara et al., 1992), and these receptors often heterodimerize and mutually regulate (Bordet et al., 2000; Marcellino et al., 2008). D2-like receptors are not abundant in NCM, but their presence cannot be ruled out (Kubikova et al., 2010). In fact, systemic D2 receptors have been shown to mediate song preference in adult female zebra finches, a phenomenon that involves NCM (Day et al., 2019). Therefore, simultaneous modulation of D1- and D2-family receptors, or of D2-family receptors alone, might be necessary to emulate dopamine effects in NCM.

We propose three models (Fig. 9B) for the effects we observed *in vivo*. If the cell types we recorded are part of the same microcircuit, it is plausible they are affecting each other’s firing properties. Therefore, in our “connected model 1”, we suggest that the D1R activation might be increasing the tonic firing of a GABAergic neuron upstream to NS1 cells, thus inhibiting them and disinhibiting BS1 cells. This model resembles a disinhibitory circuitry discovered in mammalian cortex for auditory associative learning, in which learning activates layer 1 inhibitory interneurons, which inhibit layer 2/3 PV+ interneurons, thus disinhibiting pyramidal neurons. These layer 1 neurons are activated by cholinergic signaling (Letzkus et al., 2011), and are known to be 5HT3a+/VIP– interneurons (Tremblay et al., 2016). Alternatively, our “connected model 2” depicts a single synapse and inhibitory effects of SKF-38393 on NS1 cells. Finally, our “disconnected model” summarizes our findings in each cell type and depicts isolated effects of the D1R agonist. Future experiments involving genetic targeting of specific neuronal subtypes could clarify these circuit properties.

Acetylcholine has been shown to affect stimulus-specific adaptation (SSA) in mammalian auditory cortex and inferior colliculus (Ayala and Malmierca, 2015; Metherato and Weinberger, 1989). However, to our knowledge, dopamine modulation of SSA in vertebrate auditory cortex/pallium has not been explored. In mammalian auditory cortex, D1R-induced changes in microcircuit excitability have been shown to improve signal detection in an auditory avoidance task (Happel et al., 2014), and local D1R activation improves association learning (Schicknick et al., 2012). In humans, systemic dopaminergic treatments have been shown to improve auditory language associative learning (Breitenstein et al., 2004; Knecht et al., 2004). Since dopamine signaling in auditory cortex is involved in learning, it is plausible to hypothesize that dopamine could be affecting SSA. In songbird NCM, SSA has been shown to parallel familiarity with sounds, such that novel sounds will produce more negative slopes (i.e. higher SSA) than familiar, previously adapted sounds (Chew et al., 1996). In fact, after successful behavioral association learning, learned sounds produce less SSA than novel sounds (Bell et al., 2015). Here, we provide evidence that D1Rs are involved in this process, such that pharmacological D1R activation disrupts SSA in NCM neurons. Importantly, our data show that changes in SSA do not correlate with changes in spontaneous or stimulus firing rate, suggesting that lower SSA is not the product of a firing rate “floor-effect”.

Our experiments were designed to test the prediction that blunt D1R activation would produce cellular plasticity in NCM. However, it is important to note that naturalistic dopamine signaling regulation is much more spatially and temporally targeted (Floresco et al., 2003; Grace, 1991; Mohebi et al., 2019) Therefore, it is likely that the effects observed here are a result of indiscriminate application of D1R agonist, when in reality, dopamine synaptic effects, as well as dopamine release are expected to be more nuanced spatially and temporally. With these aspects in mind, we hypothesize that indiscriminate D1R activation forces the NCM circuit into a “preadapted” state making it unable to adapt to subsequent presentation of novel sounds. Perhaps dopaminergic activation more precisely paired with sound stimuli would produce more specific changes. Therefore, future work should examine whether D1R activation in NCM paired with sounds would promote changes in SSA and association learning, including juvenile song learning.

We note that circuit origins of dopaminergic fibers to NCM are still an open question in the field. There are 8 major subpallial dopaminergic nuclei, which are fairly well conserved across vertebrates (Reiner et al., 1998). Preliminary reports suggest the caudal ventral tegmental area projects to NCM (Barr et al., 2019; Yanagihara et al., 2019), which, if confirmed, would be an interesting avenue for studying auditory reward prediction learning. Other reports suggest that the locus coeruleus (LC) projects to NCM to provide norepinephrinergic input (Chen et al., 2016; Ikeda et al., 2015). Norepinephrine is a precursor to dopamine, and LC neurons are known to release dopamine in addition to NE throughout the cortex (Devoto et al., 2005). Dopamine released by the LC onto dorsal hippocampus is involved in spatial learning and memory in rodents, independently of NE release (Kempadoo et al., 2016; Takeuchi et al., 2016). Furthermore, future studies should clarify through neuronal tract tracing which specific nuclei provide dopaminergic inputs to NCM and whether the effects observed in this study can be mimicked by dopamine release from such nuclei.

In conclusion, we show that D1R signaling shifts the excitatory-inhibitory balance in songbird pallium to modulate mechanisms involved in auditory learning and key components of auditory response, circuitry, and plasticity. We propose that D1Rs are important mediators of learning and memory in the avian sensory pallium and this mechanism could be a common feature among vertebrates.

## 1.4 Material and methods

### 1.4.1 Animals

A total of 52 adult (> 90 days old) zebra finches (34 males) were used across all experiments. Birds came from the University of Massachusetts Amherst colony (14:10 hour light-dark cycle) and were not actively breeding (single-sex cages). All procedures were in accordance with the Institutional Animal Care and Use Committee at the University of Massachusetts Amherst.

### 1.4.2 Immunofluorescence, imaging and quantification

Four males and four females were used for immunofluorescence experiments, to characterize the phenotype of D1 receptor-positive (D1R+) cells. Briefly, animals were taken from the aviary, deeply anesthetized, and perfused with ice-cold phosphate buffered saline followed by room-temperature phosphate-buffered 4% paraformaldehyde. Brains were extracted, postfixed in the same fixative overnight, cryoprotected in 30% sucrose and frozen until processing. With a cryostat, 40 µm parasagittal sections were made and sampled in 4 subseries collected in cryoprotectant solution and stored at –20 °C until processed.

Of the above subjects, tissue from 2 males and 2 females was processed for D1R, aromatase (ARO) and tyrosine hydroxylase (TH) triple immunofluorescence; tissue from all 4 males and 4 females was processed for D1R, GABA and parvalbumin (PV) triple immunofluorescence.

Sections containing NCM were selected and transferred from cryoprotectant to phosphate buffer (PB) and washed 3×15 min. They were then incubated in 10% normal goat serum (NGS; Vector Labs, Burlingame, CA) in 0.3% Triton-X (Thermo Fisher Scientific, Waltham, MA) in PB (PBT) for 2 h. Primary antibody solutions (Table 1) were prepared in 10% NGS in 0.3% PBT. To confirm antibody specificity, in a subset of sections the D1R antibody was preincubated for 1 h with blocking peptide (Fig. 1B). Sections were incubated with primary antibodies for 1 h at room temperature, followed by 2 days at 4 °C. Then, sections were washed 3×15 min in 0.1% PBT and transferred to secondary antibody solutions (goat polyclonals; Thermo Fisher; 1:200) prepared in 0.3% PBT. Finally, sections were washed in 3×10 min in 0.1% PBT and kept in the same solution in the fridge until mounted (1 or 2 days later) and coverslipped with ProLong Diamond with DAPI (Thermo Fisher).

**Table 1:** Primary antibody table

| **Antibody** | **Type** | **Host** | **Dilution** | **Company** |
| --- | --- | --- | --- | --- |
| anti-D1DR | Polyclonal | Guinea pig | 1:100 | Alomone Labs |
| D1DR peptide | - | Rat | 1:10 | Alomone Labs |
| anti-Aromatase | Polyclonal | Rabbit | 1:2000 | Azac |
| anti-Aromatase | Polyclonal | Sheep | 1:2000 | Bethyl Labs |
| anti-EGR1 | Polyclonal | Rabbit | 1:2000 | Santa Cruz |
| anti-Tyrosine Hydroxylase | Monoclonal | Mouse | 1:2000 | Immunostar |
| anti-GABA | Polyclonal | Rabbit | 1:1000 | Sigma |
| anti-Parvalbumin | Monoclonal | Mouse | 1:10000 | Millipore |

Images were taken with a confocal microscope (Nikon A1si). First, NCM was localized and a 4×4 large image was taken at 10x magnification. Then, using only the DAPI channel, the microscope stage was digitally controlled and moved to selected locations on the 10x images, at the ventral and dorsal posterior edges of NCM (Fig. 1C). Then, 15 µm (1 µm step size) z-stack images were taken at 60x magnification, starting from the top-most surface of the section. Two sections per hemisphere per animal were imaged. All laser intensities were maintained uniform across all images within experiments.

D1R antibody penetration noticeably decayed at ∼5 µm deep into the tissue, therefore only the top 5 µm of each z-stack was quantified. Cell counts were performed by a blinded experimenter using Fiji (ImageJ; NIH). Briefly, color histograms were set individually for each image so that background was predominantly dark and only strong signals were counted. Only antibody localization around the nuclei (DAPI) was included. Only cells with large, ovoidal nuclei (presumably neurons) were counted. Antibody quantification was done using the z-stack, while DAPI quantification was done using the z-max-projection image.

### 1.4.3 Classical conditioning and immediate early gene expression

#### 1.4.3.1 Stimuli and Behavioral procedure

Twenty adult zebra finches (10 males) were used in this experiment. Animals were individually isolated for ∼17.5 hours in a sound attenuation chamber. The cage set up consisted of three horizontal perches evenly spread out parallel to the cage floor (Fig. 2A). Food and water were located near the middle perch. An LCD monitor and a speaker were positioned adjacent to the cage. Behavior was recorded with a camera positioned on the opposite side of the cage from the monitor.

Animals were exposed to two Experimental conditions (Paired vs Unpaired) and two Reference conditions (Song vs Silence). The aim of the Experimental conditions was to test whether auditory-visual classical conditioning would induce EGR1 expression. Silent videos of conspecifics elicit behavioral attention and engagement and can be employed as social reinforcers for zebra finches (Galoch and Bischof, 2007). Thus, in the Paired condition (3 males, 3 females), animals were exposed to repetitions of a 2-s pure tone (2 kHz; ∼65 dB), the conditioned stimulus, followed by a 2-s delay then a 6-s silent video of a male zebra finch singing (6 different video clips were randomized), the unconditioned stimulus. Stimuli were presented 30 times (30-90 s interstimulus interval) over a 30-min window. In the Unpaired condition (3 males, 3 females), animals were presented with the same amount of stimuli (30 videos and 30 tones), but those were presented separately in pseudorandom order (15-45 s interstimulus interval) over a 30-min window. Videos were scaled so that birds in the video were displayed at approximately real sizes.Conspecific song playback after periods of isolation drive robust expression of EGR1 in NCM when compared to silence conditions (Mello and Clayton, 1994; Mello et al., 1992). Therefore, we exposed birds to Reference conditions to provide expression level references to our Experimental groups. In the Song condition, animals were exposed to three different conspecific bouts of songs (each 18-21 s long; ∼65 dB) repeated 10 times in pseudorandom order over 30 min. In the silence condition animals were not presented with any stimulus for 30 min.

#### 1.4.3.2 Immunofluorescence

EGR1 protein expression in NCM peaks between 1 and 2 h after induction (Mello and Ribeiro, 1998). Therefore, in all conditions, after the presentation of the last stimulus (or after 30 minutes in the Silence group), chamber lights were turned off (to minimize further stimulation) and animals remained in the chamber for an additional 50 minutes before they were retrieved for perfusion (∼10 min until PFA). Total time from beginning of exposure to fixative exposure was ∼1.5 h.

After perfusion, brains were processed, sectioned, and stained as described above, but with the following differences. A triple immunofluorescence protocol was performed using antibodies against NeuN, D1R, EGR1 and aromatase (Table 1) prepared in 10% donkey serum (Vector Labs) in 0.3% PBT. Donkey polyclonal secondary antibodies (Thermo Fisher; 1:200) were used. Imaging and cell counts were also performed as described above, except NeuN was used as a background stain.

#### 1.4.3.3 Behavioral scoring and analyses

Video recordings were snipped into 15-s clips around stimulus presentations using custom Python code. Only the Experimental conditions (Paired and Unpaired) were analyzed.

Clips were scored twice, once for state and once for event behaviors. State behaviors included beak direction (left, right, pointed at screen, or pointed at camera), sleeping, eating, drinking or continuously moving/flying. Event behaviors were not mutually exclusive with states, and included vocalization, singing, feather ruffling, head tilts, hopping, and gaping. Videos were scored by an experimenter blinded to the subject ID and trial order. Experimental groups could be inferred by the observer because in the Paired group, the video playback shortly followed the tone presentation. Behaviors were scored using JWatcher (Blumstein and Daniel, 2007).

Behavioral data were extracted and processed using custom Python scripts. Timestamps and continuous behavior durations were aligned to stimulus timestamps and quantified as rates (Hz; event behaviors) or percent of time spent (fraction of stimulus duration; state behaviors). Behaviors outside of the stimuli presentation windows were not analyzed. Beak direction behaviors that enabled subjects to see the screen (beak pointed left, right and towards screen) were summed to comprise a new category termed “screen-time”. Conversely, beak direction opposite to the screen (back of the head facing video), eating, drinking and sleeping were averaged to comprise a category termed “distractibility”.

### 1.4.4 *In vitro* whole-cell patch clamp

#### 1.4.4.1 Recordings

Fifteen males were used for slice recordings across two experiments. We focused these experiments on males to further explore mechanisms proposed in a previous behavioral study done in males (Macedo-Lima and Remage-Healey, 2020). We note that we did not observe systematic sex differences in the immunofluorescence and *in vivo* electrophysiology findings, but we do not discard the possibility of sex differences.

After swift decapitation, the top of the skull was resected and the head was immediately immersed in a Petri dish filled with ice-cold carbogen-aerated cutting solution (0-Mg^2+^ cutting; in mM: 222 glycerol, 25 NaHCO_3_, 2.5 KCl, 1.25 NaH_2_PO_4_, 0.5 CaCl_2_, 34 glucose, 0.4 ascorbic acid, 2 Na_2_-pyruvate, 3 myoinositol. Standard: idem except 25 glucose and 3 MgCl_2_; ∼320 mOsm/kg, pH 7.4). In the Petri dish, the cerebellum was resected and the brain was removed from the skull. Then, brain was removed from the cutting solution and placed on an ice-cold Petri dish lid covered with KimWipe (Kimberly-Clark Professional, Irving, TX). The lateral forebrain of both hemispheres was cut parasagittally to yield flat lateral surfaces, and cerebral hemispheres were bisected. The lateral edges of each hemisphere were then dabbed dry on the KimWipe, glued (cyanoacrylate) to the cutting stage and immediately immersed in the vibratome (VT1000S, Leica Biosystems, Wetzlar, Germany) chamber filled with ice-cold carbogen-aerated cutting solution. Slices were cut at 250 or 300 µm starting from the medial edge which contains NCM. NCM does not have clearly defined lateral boundaries, but song-inducible gene expression experiments indicate strong responses that extend ∼1 mm from the midline (Mello and Clayton, 1994), thus only the first three sections (∼750-900 µm) were used for recordings. After cutting, slices were transferred to 37 °C carbogen-aerated 0-Mg^2+^ or standard recording solution (0-Mg^2+^; in mM: 111 NaCl, 25 NaHCO_3_, 2.5 KCl, 1.25 NaH_2_PO_4_, 2 CaCl_2_, 28 glucose, 0.4 ascorbic acid, 2 Na-pyruvate, 3 myo-inositol; standard: idem except 25 glucose and 3 MgCl_2_; ∼320 mOsm/kg, pH 7.4). After a 30-minute recovery at 37 °C and a 30-minute stabilization at room temperature, recordings started. All recordings were performed at room temperature.

Recording pipettes (borosilicate glass) were pulled with a vertical pipette puller (PC-10, Narishige International, Tokyo, Japan) and had a tip resistance of 4-7 MΩ when submerged in the recording solution and backfilled with either K-gluconate-or CsCl-based solution, for excitatory (EPSC) and inhibitory (IPSC) postsynaptic currents recordings, respectively (*K-Gluconate-based*: in mM: ∼95 K-gluconate, 20 KCl, 0.1 CaCl_2_, 5 HEPES, 5 EGTA, 3 MgATP, 0.5 NaGTP, 20 creatine-phosphate disodium; osmolarity adjusted to ∼295 mOsm/kg with K-gluconate; pH 7.4; *CsCl-based*: in mM: ∼120 CsCl, 8 NaCl, 10 TEA-Cl, 0.2 EGTA, 10 HEPES, 2 MgATP, 0.2 NaGTP; osmolarity adjusted to ∼295 mOsm/kg with CsCl; pH 7.4). Internal solutions also contained 0.1% AlexaFluor-488-hydrazide (Thermo Fisher) and 0.1% Neurobiotin (Vector Labs) for “online” and post hoc detection (see below) of the cell, respectively.

Cells were identified with an inverted microscope (Eclipse FN1, Nikon, Tokyo, Japan) with DIC optics. Recordings were made with an EPC-10 amplifier and recorded and compensated (series resistance, slow/fast capacitance) with PatchMaster software (HEKA). Liquid junction potential was automatically subtracted. Traces were digitized at 20 KHz. Recordings were made in voltage clamp mode at -70 mV. After whole-cell configuration was achieved, cells were allowed to stabilize for 5 min. Then, baseline drug cocktails (bicuculline or DNQX) were delivered and allowed to act for a minimum of 2 min. Drug-containing solutions were gravity-delivered and flow-matched to a ∼2 mL/min peristaltic pump (Cole-Palmer MasterFlex L/S) outtake. Tissue chamber capacity was ∼1 mL. Recording quality was constantly monitored between recording blocks and recordings were aborted if series resistance compensation rose above 40 MΩ.

Nine males were used for AMPA/kainate/NMDA EPSCs (sEPSCs) recordings. For sEPSC isolation, bicuculline (20 µM; Santa Cruz) was added to the 0-Mg^2+^ recording solution. For most recordings, after bicuculline was delivered and allowed to take effect, a 1-min baseline recording was made, but for a few cells (9 out of 25), a 7-min rundown recording followed. Then, ±-SKF-38393 hydrochloride (10 or 50 µM; abcam, Cambridge, MA) (Ding and Perkel, 2002) was delivered and a 7-min recording was timed to the start of the delivery. Maximum concentration of drug is estimated to have been achieved within 1 min. Finally, a 10-min washout recording was timed to the start of SKF-38393 clearance, and complete washout is estimated to have been achieved within 1 min. Finally, to confirm the nature of the currents, D-AP-5 (50 µM; abcam, Santa Cruz, Dallas, TX or Alomone, Jerusalem, Israel) was delivered and currents were monitored for ∼5 min.

Six males were used for spontaneous GABA IPSCs (sIPSCs) recordings. For sIPSC isolation, DNQX (20 µM; Tocris) was added to the standard recording solution. Recordings were made similarly to EPSC recordings, except different cells were used for SKF (n=7) and rundown experiments (n=4). After a 1-min baseline recording, either SKF-38393 (10 µM) or nothing (rundown) was added to the recording solution and currents were monitored for 7 min. Then, a 10-min washout (or continued rundown) followed. Finally, to confirm the nature of the currents, bicuculline (20 µM) was added to the recording solution and currents were monitored for ∼5 min.

After recordings were completed, the recording pipette was slowly retrieved, and slices were drop-fixed overnight in 4% paraformaldehyde in PB. Then, they were transferred to cryoprotectant solution and kept at -20 °C until processed.

#### 1.4.4.2 Analyses

Recordings were analyzed in IgorPro 6 (WaveMetrics, Lake Oswego, OR). All traces were downsampled (5x) and lowpass filtered at 500 Hz.

In sEPSC experiments, for amplitude measurements, cells were only included (n=9) in the analysis if series resistance did not change by more than 20% from baseline values. For frequency recordings, all cells (n = 25) were analyzed, as recording quality fluctuations are not expected to interfere with their detection, due to their high amplitude (>50 pA; noise band ∼5 pA). Currents were thresholded and manually curated with NeuroMatic (Rothman and Silver, 2018). After curation, currents were automatically measured by custom IgorPro code.

For sIPSC recordings, cells whose series resistance changed more than 20% from baseline were excluded from all analyses. For each cell, one template current was manually selected and spontaneous PSCs were automatically detected using a spontaneous current detection algorithm (Clements and Bekkers, 1997) implemented by Dr. Geng-Lin Li for IgorPro. After detection, all IPSCs were measured automatically by custom IgorPro code.

For imaging cells containing neurobiotin, slices were washed 3×15 min in PB and incubated in 10% NGS in 0.3% PBT for 2 h. Sections were then incubated with rabbit anti-aromatase diluted 1:2000 in 10% NGS in 0.3% PBT for 1 h at room temperature, followed by 2 days at 4 °C. Then, sections were washed 3×15 min in 0.1% PBT and transferred to 0.3% PBT containing goat anti-aromatase at 1:200 and Streptavidin-DyLight-488 (Vector) at 1:200. Finally, sections were washed in 3×10 min in 0.1% PBT and kept in the same solution in the fridge until mounted (1 or 2 days later) and coverslipped with ProLong Diamond with DAPI (Thermo Fisher). Cells were imaged using a confocal microscope (Nikon A1si) at 20 and 100x magnification.

### 1.4.5 *In vivo* awake head-fixed electrophysiology

#### 1.4.5.1 Headpost implantation and craniotomy surgery

Five males and four females were retrieved from the single-sex cages and implanted with headposts. Briefly, under isoflurane anesthesia, custom-made headposts were lowered on the top of the beak and secured with dental cement. Skull markings over NCM (1.1 lateral, 1.4 anterior from the bifurcation of the mid-sagittal sinus, 45-degree head angle) were performed, large craniotomies over NCM were made and meninges were resected. During the recordings (described below), electrode bundles were always inserted medio-caudally to markings (within 0.5 lateral, 1.2 anterior from mid-sagittal sinus). A small craniotomy was made in an anterolateral part of the skull for the implantation of a silver ground wire using cyanoacrylate. Craniotomies were sealed with Kwik-Cast (World Precision Instruments, Sarasota, FL), and birds were allowed to recover from anesthesia. Recordings were performed within 4 days of surgery.

#### 1.4.5.2 Retrodialysis-microdrive (RetroDrive) fabrication

Custom retrodialysis probe-coupled multielectrode drives (RetroDrives) were assembled in house. RetroDrives consisted of a circular printed circuit board (PCB; Sunstone Circuits, Mulino, OR) soldered to a 36-pin connector (A79026-001, Omnetics, Minneapolis, MN), and a 5 mm 17G guide tube (stainless steel; Component Supply Company, Sparta, TN). A strand of 28G enameled copper magnet wire (Remington Industries, Johnsburg, IL) was soldered to the PCB ground. Three polyimide tubes (2x 198 µm; 1x 100 µm diameter) were inserted through the guide tube, glued side-by-side on the wall of the guide tube and cut to be flush with the guide tube. Tetrodes were made by twice-folding and twisting polyimide-coated NiCr tetrode wire (Sandvik, Stockholm, Sweden) with a tetrode spinner (LabMaker, Berlin, Germany). Tetrodes were inserted through the larger polyimide tubes (4 in each) and pinned to the PCB. A single reference wire (50 µm polyimide-coated NiCr wire; Sandvik) was inserted through the smaller polyimide tube and pinned to the PCB. Wires were glued to the top of the polyimide tubes. A microdialysis cannula (CMA8011085; Harvard Apparatus, Holliston, MA) was glued adjacent to the polyimide tubes containing tetrodes, such that a minimum of 3 mm of cannula protruded from the guide tube. Tetrodes were cut with tetrode-cutting scissors (14058-11, FST, North Vancouver, Canada) to ∼0.5 mm from the tip of the cannula. Reference wire was cut to a similar length at an acute angle. When 1 mm membrane probes (CMA8011081; Harvard Apparatus) were inserted, the tips of the probe and wires were offset by ∼0.5mm. Importantly, the horizontal distance between probe wires were ∼0.2 mm. Finally, tetrodes were gold-plated to 200-250 kΩ impedance and all wires and pins were covered with liquid electrical tape (Gardner Bender, New Berlin, WI) and allowed to dry. RetroDrives were confirmed to successfully operate in NCM using baclofen/muscimol delivery to locally silence neurons within minutes in an earlier study (Macedo-Lima and Remage-Healey, 2020).

#### 1.4.5.3 Recording protocol

On the day of the recording, a microdialysis probe was perfused with artificial cerebrospinal fluid (aCSF; described below) using a microinjection pump (PHD2000, Harvard Apparatus). RetroDrive wires were dipped in 6.25% DiI (Thermo Fisher) in 200-proof ethanol for visualization of electrode tracks. Then, the animal was comfortably restrained and head-fixed. The Kwik-Cast was removed from the craniotomy over one of the hemispheres. Animal and RetroDrive grounds were connected using alligator clips. The microdialysis probe was inserted through the cannula and the RetroDrive was lowered to NCM (∼1.5-2 mm from brain surface; mediocaudally to skull markings). Importantly, tetrodes were positioned medially to the probe such that wires were ∼0.5 mm lateral and ∼1.2 anterior from the stereotaxic zero (midsagittal sinus).

Recordings were made while animals listened to auditory stimuli and aCSF (PRE) followed by SKF-38393 (SKF) followed by aCSF (POST) were infused during song playback to assess within-subject the effects of SKF on responses to auditory stimuli (described in detail below). Recordings were completed within 4 hours of restraint.

Recordings were made from both hemispheres in different days. When recording in the first hemisphere was completed, the craniotomy was resealed with Kwik-Cast and the animal was returned to the home cage. Within 2 days, the second hemisphere recording was made, after which the animal was overdosed with isoflurane and decapitated. The brain was drop-fixed and cryoprotected in 30% sucrose in 10% formalin, and frozen until cutting. Cryostat sections were obtained at 40 µm and imaged to confirm location of wires and probe.

Recordings were amplified and digitized by a 32-channel amplifier and evaluation board (RHD2000 series; Intan Technologies, Los Angeles, CA) and sampled at 30 kHz using Intan software. An Arduino Uno (Arduino, Sommerville, MA) was connected to the recording computer to deliver TTL pulses to the evaluation board’s DAC channel bracketing the beginning and end of the audio stimuli (described below) to optimize detection during analysis. Audio playback and TTL pulses were controlled by a custom-made MatLab (MathWorks, Natick, MA) script which also controlled the Arduino and sent a copy of the audio analog signal to the evaluation board ADC channel.

#### 1.4.5.4 Stimuli

Zebra finch songs were obtained from multiple databases (http://ofer.sci.ccny.cuny.edu/song_database), therefore unlikely to have been familiar to our subjects. Twenty-four song files from unrelated birds were bandpass filtered at 0.5-15 kHz and trimmed to include two consecutive motifs without introductory notes in Adobe Audition (Adobe) and mean amplitude-normalized to 70 dB in Praat (Boersma and van Heuven, 2001). Songs were randomly and equally split into two sets, then into 3 subsets containing 4 songs each. For each animal, 1 set was used per hemisphere and, within a hemisphere recording, 1 subset was used per treatment. This was done to ensure that for each treatment birds listened to novel stimuli, because NCM neurons exhibit stimulus-specific adaptation (Chew et al., 1996). Importantly, there was no difference in neuronal firing rates to different subsets, controlled by each stimulus and within each subject and each neuron (GLM/ANOVA: Subset: χ^2^(5)=5.173, p=0.395). Therefore, responses across treatments are comparable as they were presumed to reflect responses to novel stimuli.

Each playback session consisted of 4 conspecific songs, repeated 30 times each in pseudorandomized order. Interstimulus interval was pseudorandom within the interval 5±2 s. Audio pressure was amplified to ∼65 dB as measured by a sound level meter (RadioShack, Fort Worth, TX). Playback trial duration lasted ∼20 min.

Recordings were made from each hemisphere on different days. For each animal’s first recording, the starting hemisphere was randomized in the first subjects, then counterbalanced between sexes. The stimulus set was also initially randomized, then counterbalanced across sexes and hemispheres, but the subset selected for each treatment was always randomized (www.random.org).

#### 1.4.5.5 Drugs and treatment

SKF-38393 (abcam) aliquots were made in double-distilled water (20 mM; 10 µL) (Schicknick et al., 2012) and kept at –20 °C. On the day of the experiment, one aliquot was added to 1 mL (final concentration 0.2 mM as in (Schicknick et al., 2012)) of previously frozen aCSF aliquots (in mM: 199 NaCl, 26.2 NaHCO_3_, 2.5 KCl, 1 NaH_2_PO_4_, 1.3 MgSO_4_, 2.5 CaCl_2_, 11 Glucose, 0.15 bovine serum albumin; pH 7.4). All aliquots were filtered before being loaded into the RetroDrive. Treatment and playback timeline are described in Fig 6. After each round of playback, treatment syringes were switched and a 30 min infusion period elapsed. Retrodialysis speed was set to 2 µL/min (as in Remage-Healey et al., 2010; Remage-Healey and Joshi, 2012; Vahaba et al., 2017).

#### 1.4.5.6 Analyses

Sound playback timestamps were detected using a custom-made audio convolution algorithm in MatLab.

Recordings were highpass filtered at 300 Hz and common-median filtered in MatLab (MathWorks). Single-unit sorting was done with Kilosort (Pachitariu et al., 2016). Sorting results were manually curated in Phy (https://github.com/cortex-lab/phy) and only well-isolated units (high signal-to-noise ratio; low violation of refractory period (the number of interspike intervals within a 1 ms refractory period was 0.25% median in our units); low contamination with other units; segregation in waveform PCA space) were used.

After sorting, for each single-unit, 2000 waveforms were selected pseudorandomly and measured (peak-to-peak duration and ratio; Fig 4a) in MatLab. All further data processing was done in Python and R.

Spontaneous firing rates were calculated using 500 ms preceding each stimulus playback trial. Within each treatment condition, spontaneous and stimulus firing rates were averaged across stimuli and trials. Peristimulus time histograms (PSTHs) were generated using 10-ms time bins.

Z-scores were calculated by the formula 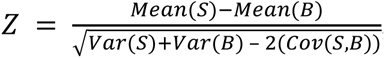, where S and B are the stimulus and spontaneous firing rates across stimulus trials, respectively. After computing z-score by stimulus, those were averaged to yield a single z-score per unit per treatment.

Adaptation rates were calculated using trials 6-25, which is the approximate-linear phase of the adaptation profile in NCM (Phan et al., 2006). For each stimulus, the stimulus firing rate across trials was normalized by the firing on trial 6 (set to 100%). Then, a linear regression was calculated between trials 6 and 25. For each treatment, the minimum (steepest) adaptation slope across stimuli was used for each unit.

Latency to respond to stimuli were calculated as in Ono et al. (2016). Briefly, for each stimulus, 5-ms PSTH were generated and convolved with a 5-point box-filter. The latency to respond to a stimulus was the time after stimulus onset in which the filtered PSTH rose above 3 standard deviations of the average preceding spontaneous firing period (100 ms). If threshold was not crossed within 400 ms, that stimulus was excluded from analyses.

### 1.4.6 Statistical approach

All statistical analyses and plotting were performed using libraries for R and Python, respectively.

Our general statistical approach was to perform generalized linear modeling (GLM; using ‘lme4’, ‘glmmTMB’, ‘lmerTest’, ‘DHARMa’ and ‘car’ R packages) followed by ANOVA. Data were initially fitted with gaussian distributions. Normality of residuals was assessed using DHARMa and Q-Q plot inspection following the GLM fits. If residuals violated normality, data were refit with other distributions (Poisson or negative binomial) and residuals were reassessed based on new distributions. If normality was still violated, data were log-transformed when possible (non-negative, non-zero data) or rankit-transformed (Bliss et al., 1956) and the fitting process above was repeated. If residuals were distributed according to GLM distributions, the model was analyzed by ANOVA using Wald chi-square tests. If data still violated residual diagnostics, non-parametric one-way analyses were performed (e.g. Kruskal-Wallis followed by Dunn’s post-hoc tests, Friedman test or Wilcoxon signed rank tests).

For immunofluorescence data, quantifications of the two sections belonging to the same hemisphere and animal were averaged. Response variables were always percentages of the total number of cells (DAPI) or neurons (NeuN). Data were analyzed by GLM followed by ANOVA, with Hemisphere and Area (dorsal vs ventral NCM) as interacting fixed factors, controlled for Area nested within Hemisphere nested within Subject. In the GABA-D1R-PV analyses, tissue from the right hemisphere of one female was excluded because NCM could not be confidently localized (off-plane section). In the EGR1-D1R-ARO analyses, data from one male in the Paired condition could not be quantified because of bad tissue quality after processing. Although animals of both sexes were used, we did not have power to detect sex differences. Nevertheless, no qualitative sex differences were observed. For EGR1 induction analyses, GLM followed by ANOVA was used, with Hemisphere, Area and Condition (Song vs Silence; Paired vs Unpaired) as interacting fixed factors, controlled for Area nested within Hemisphere nested within Subject. To test coexpression proportions, chi-square tests were performed on the total sum of cells across subjects, hemispheres and areas to form a 2×2 contingency table (e.g. ARO+/ARO– vs D1R+/D1R–). Pearson’s z-scored residuals were analyzed to obtain corresponding one-tailed p-values.

For behavioral analyses, data were grouped by stimulus type (audio or video) and averaged across all 30 trials. Then, we ran GLM followed by ANOVA, with Condition (Paired vs Unpaired) as fixed factor, controlled by Subject.

To evaluate whether behaviors predicted EGR1 expression we fit GLMs including Hemisphere, Area and Condition (Paired vs Unpaired) as interacting fixed factors and added the behavioral measurement (e.g. screen-time) as an interacting covariate, controlled for Area nested within Hemisphere nested within Subject. To test the effects of behavior on EGR1 expression with increased statistical power, for every combination of the hemisphere*area interaction we ran individual GLM/ANOVA with Condition and behavioral measurement as interacting factor and covariate.

For patch clamp recordings, minute-by-minute-binned data were analyzed by GLM followed by ANOVA, using Time bins (60 seconds) as a fixed factor controlled over Cell nested within Subject, and Treatment or Dose as independent fixed factor. Bin 1 corresponded to the minute before SKF treatment; bins 2-8 corresponded to the SKF treatment. Washout was excluded from these analyses and is presented for visualization purposes in all figures. Post-hoc Dunnett tests were used to compare treatment bins with the pre-drug bin (control).

For *in vivo* recordings, cell types were classified using 2-D hierarchical clustering (‘stats’ R package) on peak-to-peak duration vs ratio measurements. The optimal number of clusters was determined using the package Nbclust (Charrad et al., 2014) with the gap statistic method. After clustering, each unit’s auditory responsiveness was tested by Wilcoxon tests (30x spontaneous vs 30x stimulus trials per song during aCSF infusion). Cells responsive to at least one song were included in all following analyses. Before-drug (PRE) differences among cell types were tested using GLM/ANOVA (controlled over Subject) or Kruskal-Wallis tests and Dunn’s post-hoc tests with Benjamini-Yekutieli false-discovery rate adjustments. Treatment data were analyzed by GLM followed by ANOVA using Treatment and Cell type as interacting fixed factors, controlled for Unit nested within Subject; Tukey post-hoc tests were used when ANOVA effects with more than 2 factors were significant. To appropriately fit negative-binomial GLMs, firing rate data was summed (instead of averaged) across the 30 stimulus presentations. For a similar reason, to fit gaussian GLMs, z-score data were offset to not contain zero-values [*z*scores + 2 × min(*z*scores)] and log-transformed. For simplicity, average firing rates and untransformed z-scores are plotted.

Due to statistical power limitations, we performed separate analyses excluding Cell type and including Hemisphere and Sex as factors. However, no systematic sex or hemisphere differences with treatment were detected. Washout (POST) was excluded from all analyses.

## Acknowledgments

We thank current and former members of the Healey Lab at the University of Massachusetts who helped with this project, especially Amanda Krentzel, Catherine de-Bournonville, Christina Moschetto, Daniel Pollak, Daniel Vahaba, Garrett Scarpa, Jeremy Spool, Katie Schroeder, Maaya Ikeda, Marcela Fernandez-Peters and Rachel Frazier. Finally, we thank Geng-Lin Li, Joseph Bergan, Jeffrey Podos and Karine Fenelon for valuable input on this project.

---

**Supplementary figure 1.**
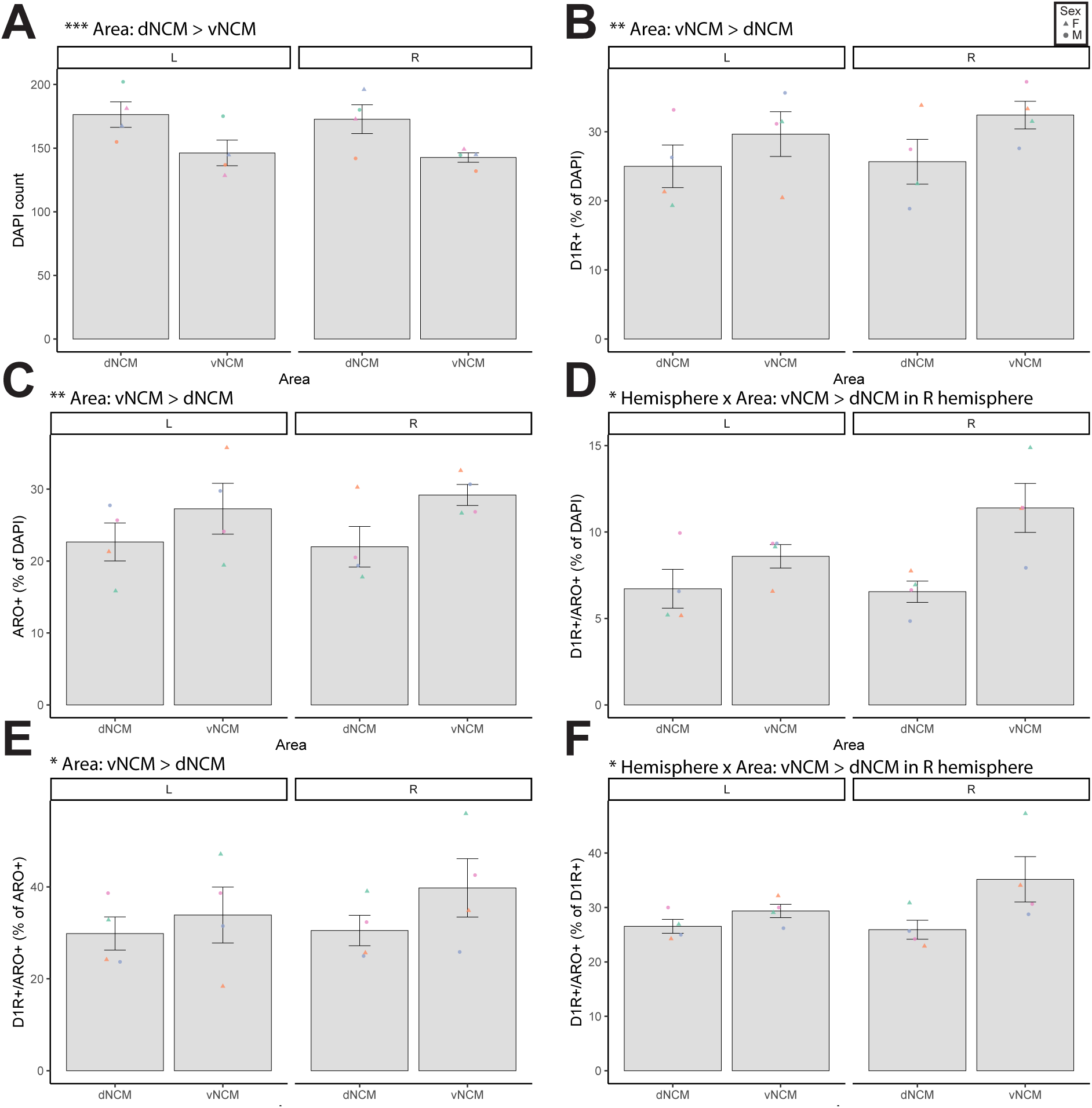
Anatomical distribution of D1 dopamine receptor+ (D1R+) and aromatase+ (ARO+) neurons in NCM. (A) DAPI counts were higher in dNCM than in vNCM. (B) Frequency of D1R+ (% of DAPI) cells was higher in vNCM than dNCM. (C) Frequency of ARO+ (% of DAPI) cells was higher in vNCM than in dNCM. (D) Frequency of D1R+/ARO+ (% of DAPI) cells was higher in the vNCM of the R hemisphere. (E) Frequency of D1R+/ARO+ (% of ARO+) cells was higher in vNCM than in dNCM. (F) Frequency of D1R+/ARO+ (% of D1R+) cells was higher in vNCM than in dNCM of the R hemisphere. Statistical results are from GLM/ANOVA (Hemisphere*Area controlled by Subject) followed by Tukey’s post-hoc tests. *p<0.05; **p<0.01; ***p<0.001. Marker colors represent data from individual subjects.

**Supplementary figure 2.**
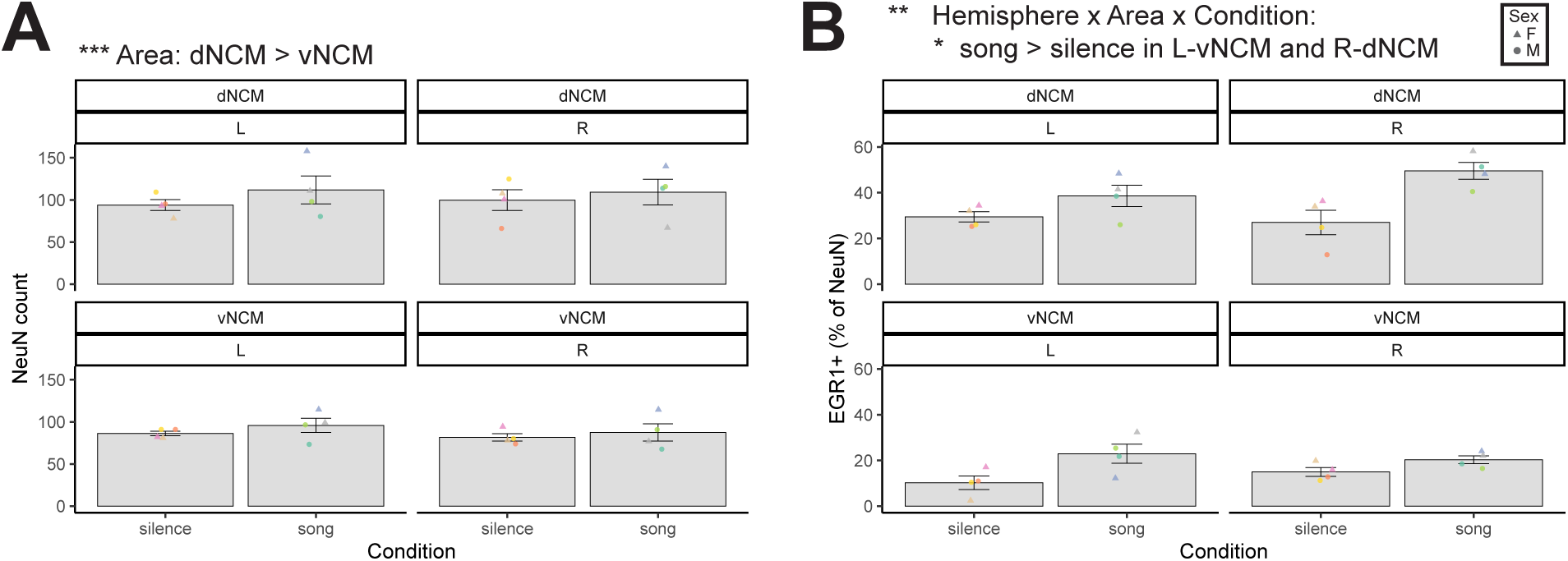
Distribution of NeuN and EGR1+ following conspecific song/silence exposure. (A) NeuN counts were higher in dNCM than in vNCM. (B) Frequency of EGR1+ (% of NeuN) neurons was higher in the Song condition, particularly in L-vNCM and R-dNCM. Statistical results are from GLM/ANOVA (Hemisphere*Area*Condition controlled by Subject) followed by Tukey’s post-hoc tests. *p<0.05; **p<0.01; ***p<0.001. Marker colors represent data from individual subjects.

**Supplementary figure 3.**
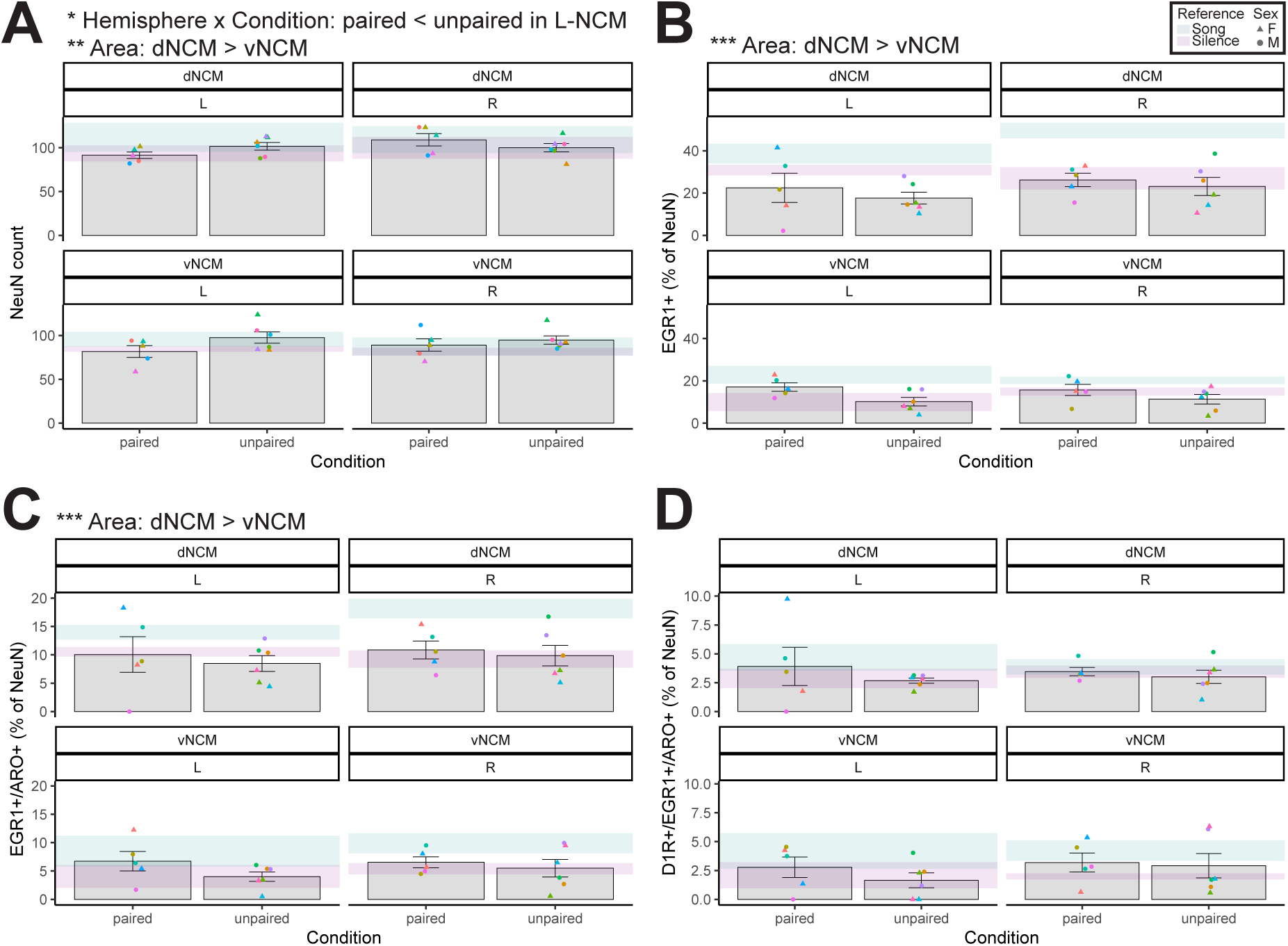
Distribution of NeuN+, EGR1+, dopamine D1 receptor+ (D1R+) and aromatase+ (ARO+) neurons in NCM following classical conditioning paradigm (Paired) or unconditioned stimulation (Unpaired). (A) NeuN expression was higher in L-NCM of the Unpaired condition and higher in dNCM than in vNCM. (B) Frequency of EGR1+ (% of NeuN) neurons was higher in dNCM than in vNCM. (C) Frequency of EGR1+/ARO+ (% of NeuN) neurons was higher in dNCM than in vNCM. (D) Frequency of D1R+/EGR1+/ARO+ (% of NeuN) neurons did not differ. Statistical results are from GLM/ANOVA (Hemisphere*Area*Condition controlled by Subject) followed by Tukey’s post-hoc tests. *p<0.05; **p<0.01; ***p<0.001. Marker colors represent data from individual subjects.

**Supplementary figure 4.**
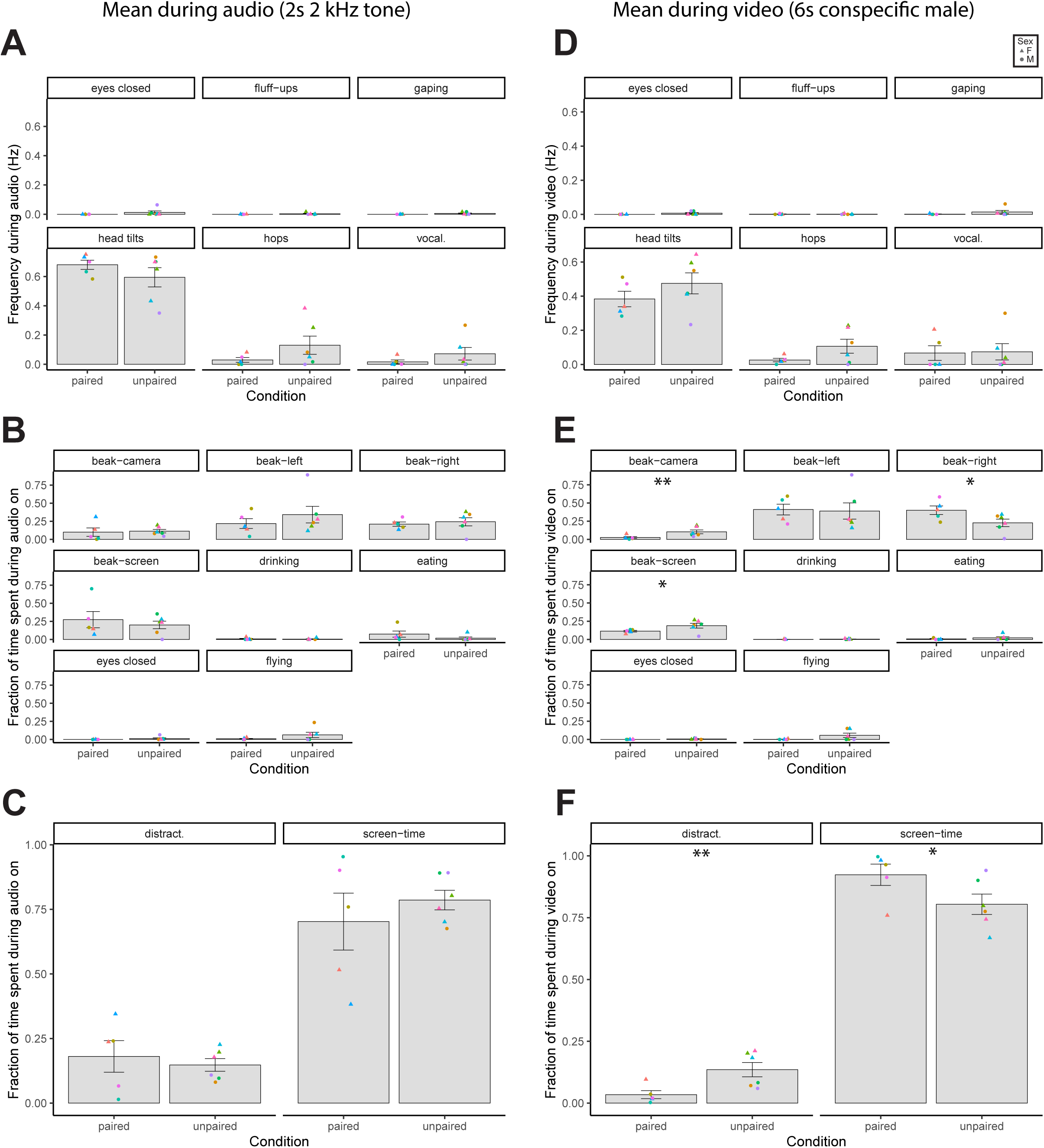
Behavioral measurements during classical conditioning paradigm (Paired) or unconditioned stimulation (Unpaired). Values were averaged across the 30 trials during audio (left) and video (right) presentations. (A) Event behaviors, (B) state behaviors, (C) distractibility (beak-camera + drinking + eating + eyes closed + flying) and screen-time (beak-left + beak-right + beak-screen) behaviors during audio presentation did not differ between groups. (D) State behaviors during video presentations did not differ between groups. (E) Beak towards right wall (i.e. left eye towards video) was higher in Paired than Unpaired condition. Beak towards camera and screen were higher in the unpaired condi-tion. (F) Distractibility and Screen-time behaviors were higher in Unpaired and in Paired conditions, respectively. Statisti-cal results are from GLM/ANOVA (Condition controlled by Subject). *p<0.05; **p<0.01; ***p<0.001. Marker colors represent data from individual subjects.

**Supplementary figure 5.**
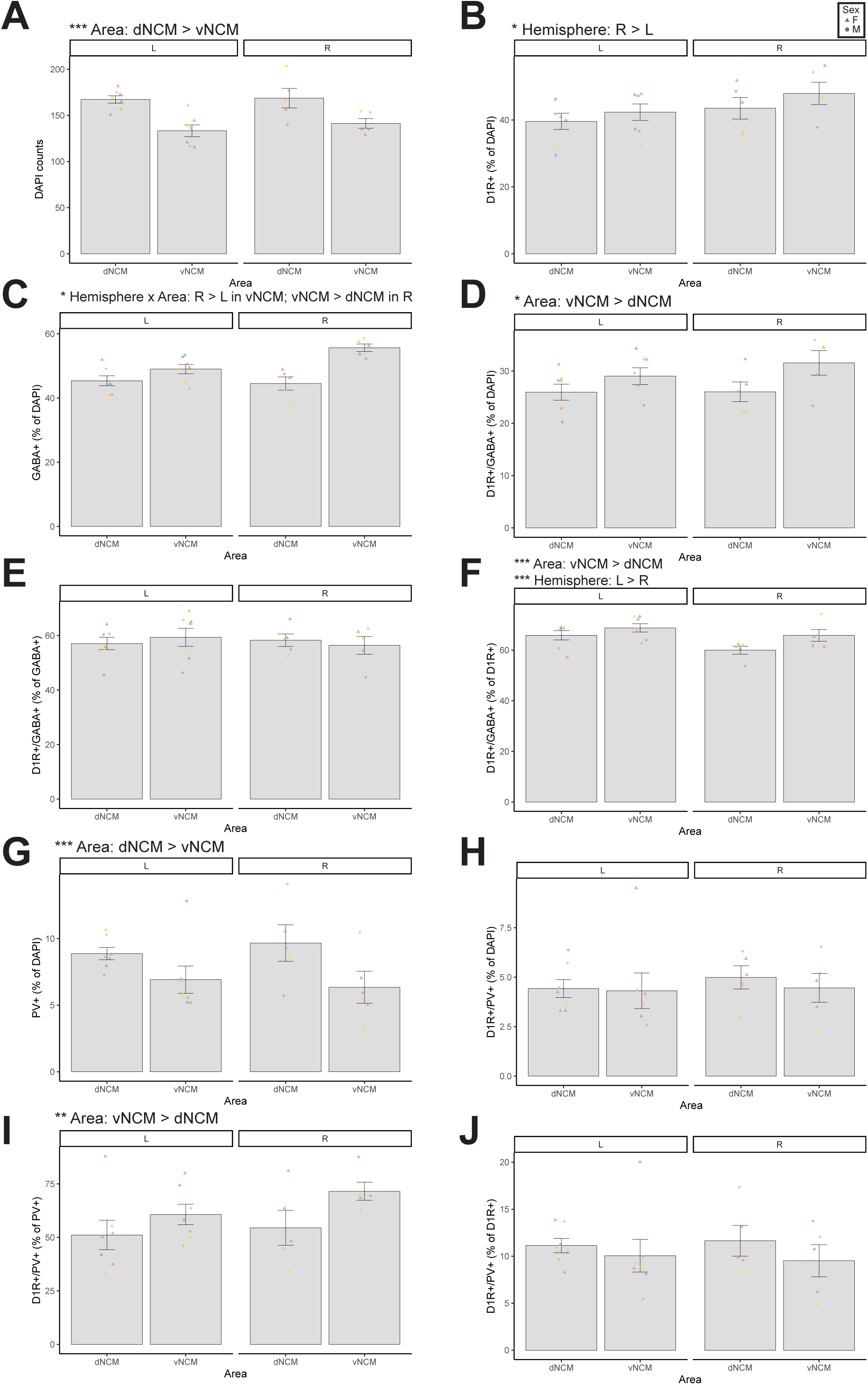
Anatomical distribution of D1 dopamine receptor+ (D1R+), GABA+ and parvalbumin+ (PV+) neurons in NCM.(A) DAPI counts were higher in dNCM than in vNCM. (B) Frequency of D1R+ (% of DAPI) neurons was higher in R than L hemisphere. (C) Frequency of GABA+ (% of DAPI) neurons was higher in the R than L hemisphere for vNCM and higher in vNCM than dNCM in the R hemisphere. (D) Frequency of D1R+/GABA+ (% of DAPI) was higher in vNCM than dNCM. (E) Frequency of D1R+/GABA+ (% of GABA+) did not differ. (F) Frequency of D1R+/GABA+ (% of D1R+) was higher in vNCM than dNCM and in L than R hemisphere. (G) Frequency of PV+ (% of DAPI) was higher in dNCM than in vNCM (H) Frequency of D1R+/PV+ (% of DAPI) did not differ. (I) Frequency of D1R+/PV+ (% of PV+) was higher in vNCM than in dNCM. (J) Frequency of D1R+/PV+ (% of D1R+) did not differ. Statistical results are from GLM/ANOVA (Hemisphere*Area controlled by Subject) followed by Tukey’s post-hoc tests. *p<0.05; **p<0.01; ***p<0.001. Marker colors represent data from individual subjects.

**Supplementary figure 6.**
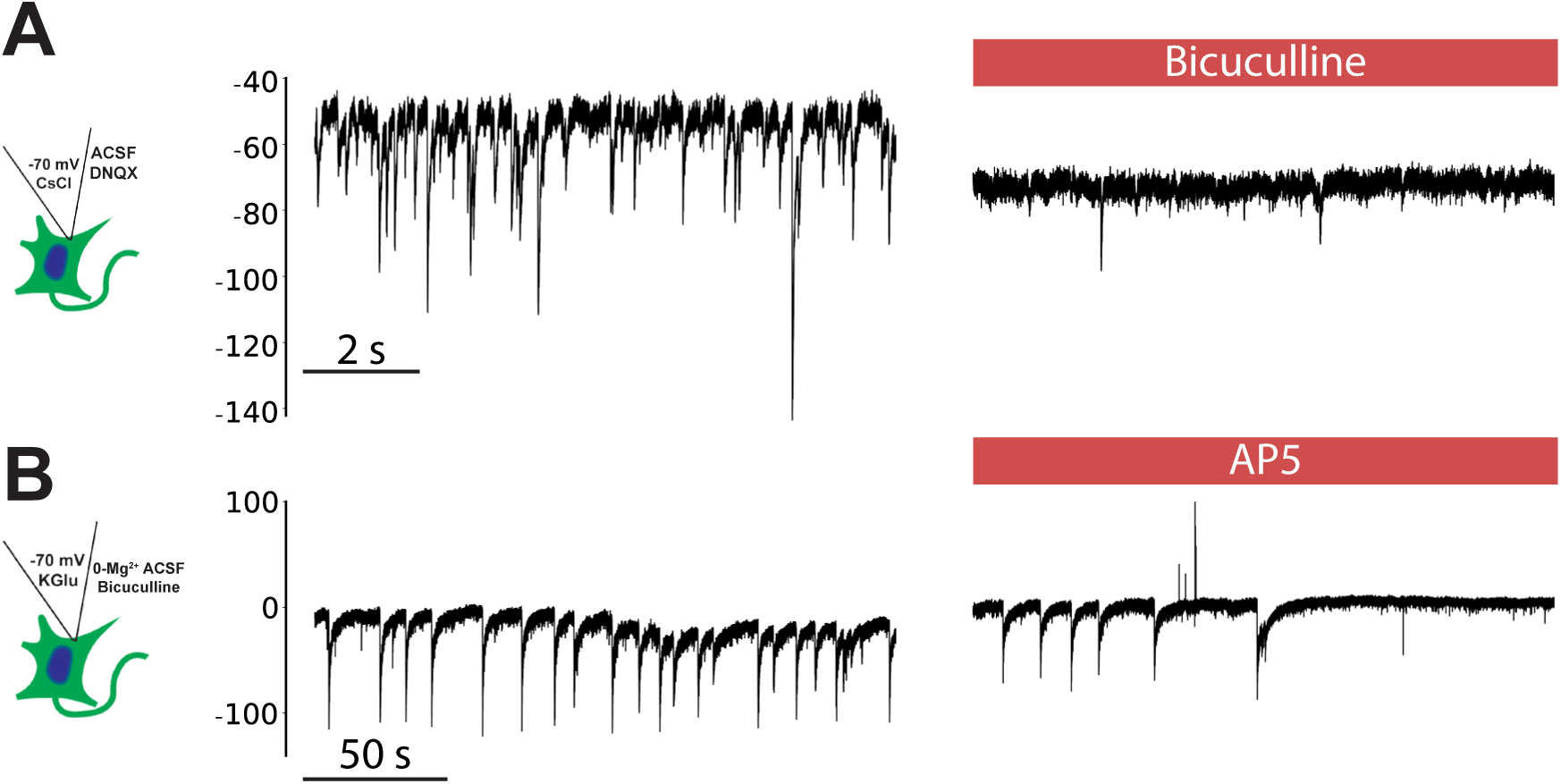
Pharmacological confirmation of spontaneous currents for whole-cell in vitro electrophysiolo-gy. (A) Spontaneous GABAergic currents were inhibited with bicuculline at the end of recordings. (B) Spontaneous NMDA-dependent glutamatergic currents were inhibited by AP5 at the end of the recordings.

**Supplementary figure 7.**
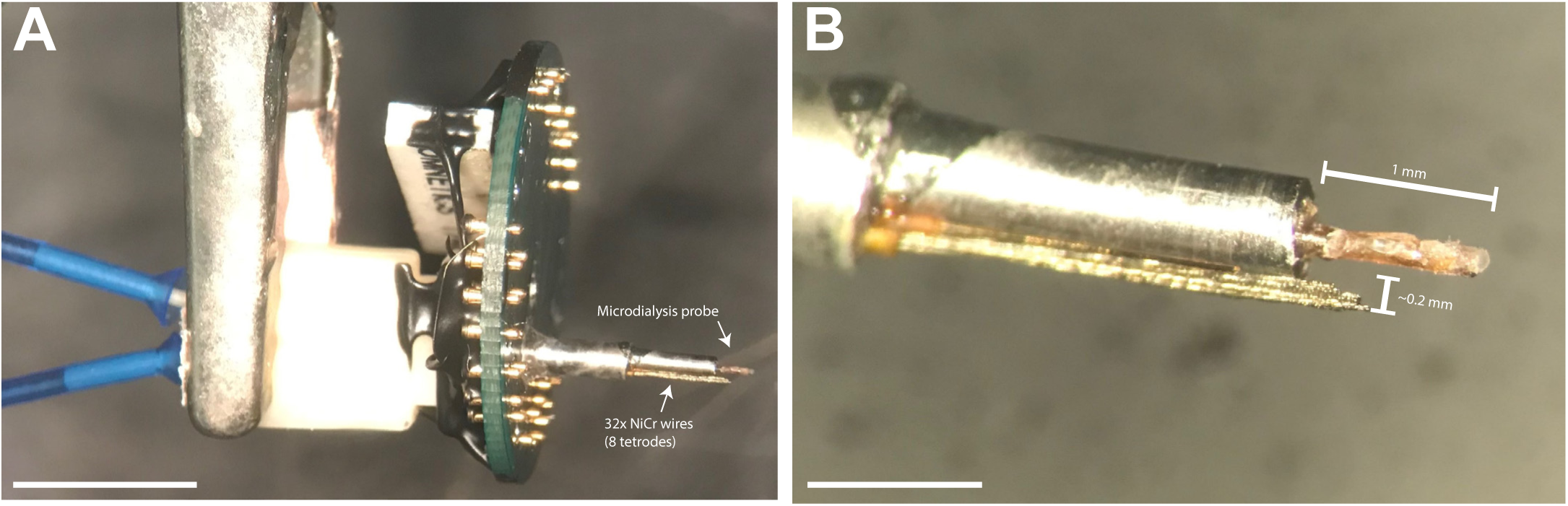
RetroDrive: acute multichannel electrophysiology microdrive coupled to a microdialysis probe for drug delivery through retrodialysis. (A) RetroDrives were constructed using a custom electrode interface board, a 32-channel Omnetics connector, a CMA11 (Harvard Apparatus) microdialysis probe, and 32x 12.5-µm-diameter NiCr wires spun into tetrodes. Scale bar: 5 mm. (B) Tetrodes were ∼0.2 mm distant from the probe and cut to extend to approxi-mately half the length of the probe. Scale bar: 1 mm.

**Supplementary figure 8.**
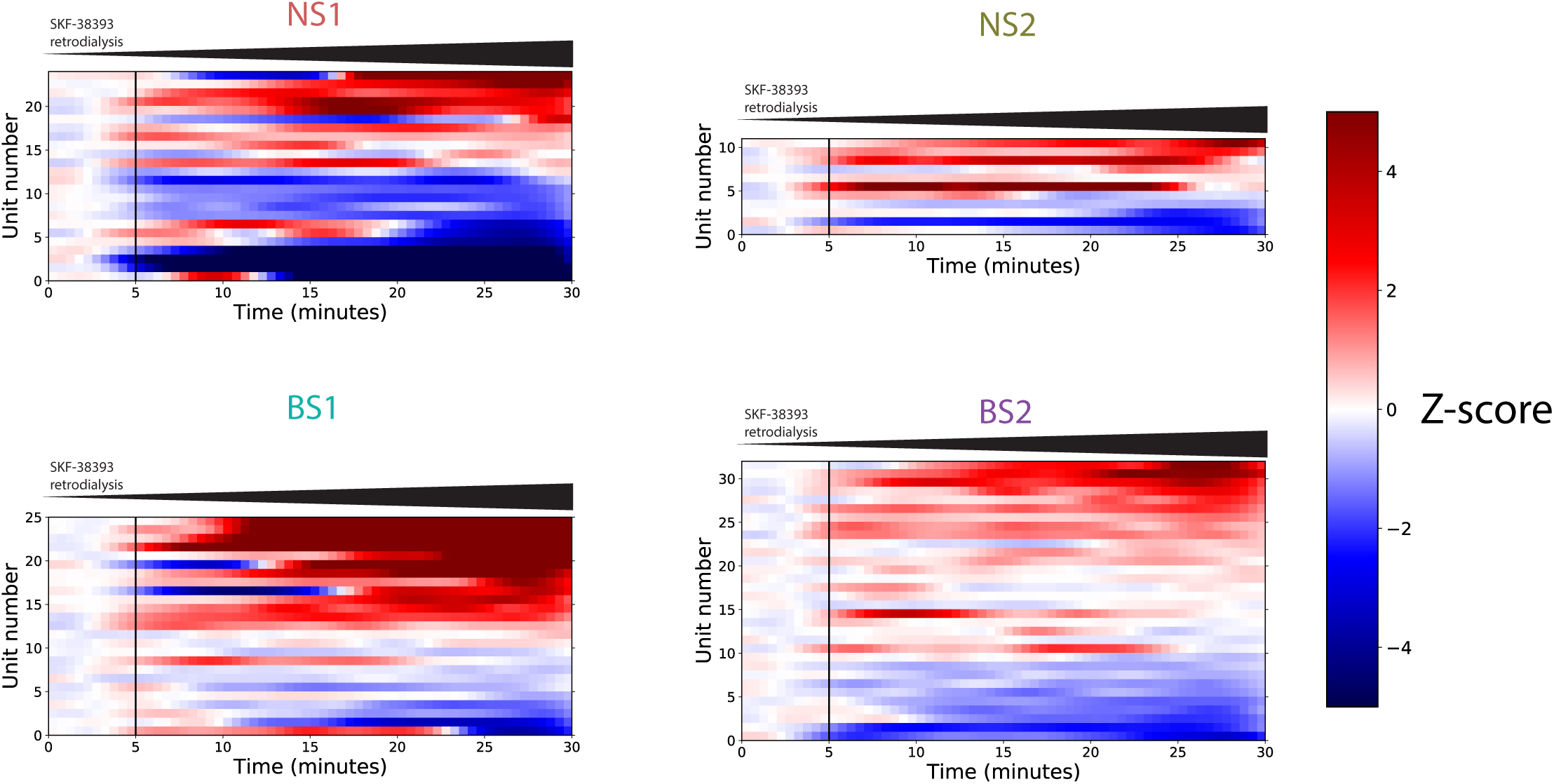
Spontaneous activity (z-scored) of single units in awake extracellular recordings during SKF-38393 (D1 receptor agonist) infusion. The start of the microdialysis pump is at time=0. We estimate the drug reached the probe at ∼5 minutes. Activity was 30-s-binned and z-score-normalized by the average spontaneous firing during the first 5 minutes. Heatmaps were ordered in decreasing order (mean of last 5 minutes). Cell-type classifications (NS1, NS2, BS1 and BS2) are based on unsupervised hierarchical clustering (Fig. 5)

## References

Atallah B V., Bruns W, Carandini M, Scanziani M. 2012. Parvalbumin-Expressing Interneurons Linearly Transform Cortical Responses to Visual Stimuli. Neuron 73:159–170. doi:10.1016/j.neuron.2011.12.013

Atencio CA, Schreiner CE. 2008. Spectrotemporal processing differences between auditory cortical fast-spiking and regular-spiking neurons. J Neurosci 28:3897–3910. doi:10.1523/JNEUROSCI.5366-07.2008

Aurore C, Giret N, Edeline J-M, Del Negro C. 2019. Neuronal encoding in a high level auditory area: from sequential order of elements to grammatical structure. J Neurosci 2767–18. doi:10.1523/JNEUROSCI.2767-18.2019

Ayala YA, Malmierca MS. 2015. Cholinergic modulation of stimulus-specific adaptation in the inferior colliculus. J Neurosci 35:12261–12272. doi:10.1523/JNEUROSCI.0909-15.2015

Balmer TS, Carels VM, Frisch JL, Nick TA. 2009. Modulation of perineuronal nets and parvalbumin with developmental song learning. J Neurosci 29:12878–12885. doi:10.1523/JNEUROSCI.2974-09.2009

Balthazart J, Baillien M, Ball GF. 2002. Interactions between aromatase (estrogen synthase) and dopamine in the control of male sexual behavior in quail. Comp Biochem Physiol - B Biochem Mol Biol 132:37–55. doi:10.1016/S1096-4959(01)00531-0

Bao S, Chan VT, Merzenich MM. 2001. Cortical remodelling induced by activity of ventral tegmental dopamine neurons. Nature 412:79–83. doi:10.1038/35083586

Barr HJ, Wall EM, Woolley SC. 2019. Dopamine in the songbird auditory cortex shapes auditory preference. bioRxiv 761783. doi:10.1101/761783

Beaulieu J-M, Gainetdinov RR. 2011. The physiology, signaling, and pharmacology of dopamine receptors. Pharmacol Rev 63:182–217. doi:10.1124/pr.110.002642.182

Becker JB. 1990. Estrogen rapidly potentiates amphetamine-induced striatal dopamine release and rotational behavior during microdialysis. Neurosci Lett 118:169–171. doi:10.1016/0304-3940(90)90618-J

Bell BA, Phan ML, Vicario DS. 2015. Neural responses in songbird forebrain reflect learning rates, acquired salience, and stimulus novelty after auditory discrimination training. J Neurophysiol 113:1480–92. doi:10.1152/jn.00611.2014

Bliss CI, Greenwood ML, White ES. 1956. A Rankit Analysis of Paired Comparisons for Measuring the Effect of Sprays on Flavor. Biometrics 12:381. doi:10.2307/3001679

Blumstein DT, Daniel JC. 2007. Quantifying behavior the JWatcher way. Sunderland (MA): Sinauer Associates Incorporated.

Boersma P, van Heuven V. 2001. Speak and unSpeak with Praat. Glot Int 5:341–347.

Bolhuis JJ, Okanoya K, Scharff C. 2010. Twitter evolution: converging mechanisms in birdsong and human speech. Nat Rev Neurosci 11:747–59. doi:10.1038/nrn2931

Bordet R, Ridray S, Schwartz JC, Sokoloff P. 2000. Involvement of the direct striatonigral pathway in levodopa-induced sensitization in 6-hydroxydopamine-lesioned rats. Eur J Neurosci 12:2117–2123. doi:10.1046/j.1460-9568.2000.00089.x

Bottjer SW, Ronald AA, Kaye T. 2019. Response properties of single neurons in higher level auditory cortex of adult songbirds. J Neurophysiol 121:218–237. doi:10.1152/jn.00751.2018

Breitenstein C, Wailke S, Bushuven S, Kamping S, Zwitserlood P, Ringelstein EB, Knecht S. 2004. D-amphetamine boosts language learning independent of its cardiovascular and motor arousing effects. Neuropsychopharmacology 29:1704–1714. doi:10.1038/sj.npp.1300464

Calabresi P, Maj R, Mercuri NB, Bernardi G. 1992. Coactivation of D1 and D2 dopamine receptors is required for long-term synaptic depression in the striatum. Neurosci Lett 142:95–99. doi:10.1016/0304-3940(92)90628-K

Cardin JA. 2018. Inhibitory Interneurons Regulate Temporal Precision and Correlations in Cortical Circuits. Trends Neurosci 41:689–700. doi:10.1016/j.tins.2018.07.015

Charrad M, Ghazzali N, Boiteau V, Niknafs A. 2014. Nbclust: An R package for determining the relevant number of clusters in a data set. J Stat Softw 61:1–36. doi:10.18637/jss.v061.i06

Chen Y, Matheson LE, Sakata JT. 2016. Mechanisms underlying the social enhancement of vocal learning in songbirds. Proc Natl Acad Sci 201522306. doi:10.1073/pnas.1522306113

Chew SJ, Vicario DS, Nottebohm FN. 1996. A large-capacity memory system that recognizes the calls and songs of individual birds. Proc Natl Acad Sci U S A 93:1950–5. doi:10.1073/pnas.93.5.1950

Clements JD, Bekkers JM. 1997. Detection of spontaneous synaptic events with an optimally scaled template. Biophys J 73:220–229. doi:10.1016/S0006-3495(97)78062-7

Darvish-Ghane S, Yamanaka M, Zhuo M. 2016. Dopaminergic Modulation of Excitatory Transmission in the Anterior Cingulate Cortex of Adult Mice. Mol Pain 12:1–14. doi:10.1177/1744806916648153

Day NF, Saxon D, Robbins A, Harris L, Nee E, Shroff-Mehta N, Stout K, Sun J, Lillie N, Burns M, Korn C, Coleman MJ. 2019. D2 dopamine receptor activation induces female preference for male song in the monogamous zebra finch. J Exp Biol 222:jeb191510. doi:10.1242/jeb.191510

Devoto P, Flore G, Saba P, Fà M, Gessa GL. 2005. Co-release of noradrenaline and dopamine in the cerebral cortex elicited by single train and repeated train stimulation of the locus coeruleus. BMC Neurosci 6:1–11. doi:10.1186/1471-2202-6-31

Ding L, Perkel DJ. 2002. Dopamine Modulates Excitability of Spiny Neurons in the Avian Basal Ganglia. J Neurosci 22:5210–5218. doi:10.1523/jneurosci.22-12-05210.2002

Donato F, Rompani SB, Caroni P. 2013. Parvalbumin-expressing basket-cell network plasticity induced by experience regulates adult learning. Nature 504:272–276. doi:10.1038/nature12866

Duclot F, Kabbaj M. 2017. The role of early growth response 1 (EGR1) in brain plasticity and neuropsychiatric disorders. Front Behav Neurosci 11:1–20. doi:10.3389/fnbeh.2017.00035

Floresco SB, West AR, Ash B, Moorel H, Grace AA. 2003. Afferent modulation of dopamine neuron firing differentially regulates tonic and phasic dopamine transmission. Nat Neurosci 6:968–973. doi:10.1038/nn1103

Gadagkar V, Puzerey PA, Chen R, Baird-Daniel E, Farhang AR, Goldberg JH. 2016. Dopamine neurons encode performance error in singing birds. Science (80-) 354. doi:10.1126/science.aah6837

Galoch Z, Bischof HJ. 2007. Behavioural responses to video playbacks by zebra finch males. Behav Processes 74:21–26. doi:10.1016/j.beproc.2006.09.002

Gonzalez-Islas C, Hablitz JJ. 2003. Dopamine enhances EPSCs in layer II-III pyramidal neurons in rat prefrontal cortex. J Neurosci 23:867–875. doi:10.1523/jneurosci.23-03-00867.2003

Grace AA. 1991. Phasic versus tonic dopamine release and the modulation of dopamine system responsivity: A hypothesis for the etiology of schizophrenia. Neuroscience 41:1–24. doi:10.1016/0306-4522(91)90196-U

Happel MFK, Deliano M, Handschuh J, Ohl FW. 2014. Dopamine-Modulated Recurrent Corticoefferent Feedback in Primary Sensory Cortex Promotes Detection of Behaviorally Relevant Stimuli. J Neurosci 34:1234–1247. doi:10.1523/JNEUROSCI.1990-13.2014

Hromádka T, DeWeese MR, Zador AM. 2008. Sparse Representation of Sounds in the Unanesthetized Auditory Cortex. PLoS Biol 6:e16. doi:10.1371/journal.pbio.0060016

Ichihara K, Nabeshima T, Kameyama T. 1992. Effects of dopamine receptor agonists on passive avoidance learning in mice: interaction of dopamine D1 and D2 receptors. Eur J Pharmacol 213:243–249. doi:10.1016/0014-2999(92)90688-Z

Ikeda MZ, Jeon SD, Cowell R a., Remage-Healey L. 2015. Norepinephrine Modulates Coding of Complex Vocalizations in the Songbird Auditory Cortex Independent of Local Neuroestrogen Synthesis. J Neurosci 35:9356–9368. doi:10.1523/JNEUROSCI.4445-14.2015

Ikeda MZ, Krentzel AA, Oliver TJ, Scarpa GB, Remage-Healey L. 2017. Clustered organization and region-specific identities of estrogen-producing neurons in the forebrain of Zebra Finches (Taeniopygia guttata). J Comp Neurol 525:3636–3652. doi:10.1002/cne.24292

Jarvis ED. 2019. Evolution of vocal learning and spoken language. Science (80-) 366:50–54. doi:10.1126/science.aax0287

Kempadoo KA, Mosharov E V., Choi SJ, Sulzer D, Kandel ER. 2016. Dopamine release from the locus coeruleus to the dorsal hippocampus promotes spatial learning and memory. Proc Natl Acad Sci U S A 113:14835–14840. doi:10.1073/pnas.1616515114

Knecht S, Breitenstein C, Bushuven S, Wailke S, Kamping S, Floel A, Zwitserlood P, Ringelstein EB. 2004. Levodopa: Faster and better word learning in normal humans. Ann Neurol 56:20–26. doi:10.1002/ana.20125

Krentzel AA, Ikeda MZ, Oliver TJ, Koroveshi E, Remage-Healey L. 2019. Acute neuroestrogen blockade attenuates song-induced immediate early gene expression in auditory regions of male and female zebra finches. J Comp Physiol A 206:15–31. doi:10.1007/s00359-019-01382-w

Kubikova L, Wada K, Jarvis ED. 2010. Dopamine receptors in a songbird brain. J Comp Neurol 518:741–769. doi:10.1002/cne.22255

Lammers CH, D’Souza U, Qin ZH, Lee SH, Yajima S, Mouradian MM. 1999. Regulation of striatal dopamine receptors by estrogen. Synapse 34:222–7. doi:10.1002/(SICI)1098-2396(19991201)34:3<222::AID-SYN6>3.0.CO;2-J

Lampen J, McAuley JD, Chang SE, Wade J. 2017. ZENK induction in the zebra finch brain by song: Relationship to hemisphere, rhythm, oestradiol and sex. J Neuroendocrinol 29:1–10. doi:10.1111/jne.12543

Le Moine C, Gaspar P. 1998. Subpopulations of cortical GABAergic interneurons differ by their expression of D1 and D2 dopamine receptor subtypes. Mol Brain Res 58:231–236. doi:10.1016/S0169-328X(98)00118-1

Leblois A, Wendel BJ, Perkel DJ. 2010. Striatal Dopamine Modulates Basal Ganglia Output and Regulates Social Context-Dependent Behavioral Variability through D1 Receptors. J Neurosci 30:5730–5743. doi:10.1523/JNEUROSCI.5974-09.2010

Lee V, Pawlisch BA, Macedo-Lima M, Remage-Healey L. 2018. Norepinephrine enhances song responsiveness and encoding in the auditory forebrain of male zebra finches. J Neurophysiol 119:209–220. doi:10.1152/jn.00251.2017

Letzkus JJ, Wolff SBE, Meyer EMM, Tovote P, Courtin J, Herry C, Lüthi A. 2011. A disinhibitory microcircuit for associative fear learning in the auditory cortex. Nature 480:331–335. doi:10.1038/nature10674

Lu K, Vicario DS. 2017. Familiar but unexpected: Effects of sound context statistics on auditory responses in the songbird forebrain. J Neurosci 37:12006–12017. doi:10.1523/JNEUROSCI.5722-12.2017

Lüscher C, Malenka RC. 2012. NMDA receptor-dependent long-term potentiation and long-term depression (LTP/LTD). Cold Spring Harb Perspect Biol 4:1–16. doi:10.1101/cshperspect.a005710

Macedo-Lima M, Remage-Healey L. 2020. Auditory learning in an operant task with social reinforcement is dependent on neuroestrogen synthesis in the male songbird auditory cortex. Horm Behav 121:104713. doi:10.1016/j.yhbeh.2020.104713

Maney DL, Cho E, Goode CT. 2006. Estrogen-dependent selectivity of genomic responses to birdsong. Eur J Neurosci 23:1523–1529. doi:10.1111/j.1460-9568.2006.04673.x

Marcellino D, Ferré S, Casadó V, Cortés A, Le Foll B, Mazzola C, Drago F, Saur O, Stark H, Soriano A, Barnes C, Goldberg SR, Lluis C, Fuxe K, Franco R. 2008. Identification of dopamine D1-D3 receptor heteromers: Indications for a role of synergistic D1-D3 receptor interactions in the striatum. J Biol Chem 283:26016–26025. doi:10.1074/jbc.M710349200

Matheson LE, Sakata JT. 2015. Catecholaminergic contributions to vocal communication signals. Eur J Neurosci n/a-n/a. doi:10.1111/ejn.12885

Matragrano LL, Beaulieu M, Phillip JO, Rae AI, Sanford SE, Sockman KW, Maney DL. 2012. Rapid effects of hearing song on catecholaminergic activity in the songbird auditory pathway. PLoS One 7:e39388. doi:10.1371/journal.pone.0039388

Matragrano LL, Sanford SE, Salvante KG, Sockman KW, Maney DL. 2011. Estradiol-dependent catecholaminergic innervation of auditory areas in a seasonally breeding songbird. Eur J Neurosci 34:416–25. doi:10.1111/j.1460-9568.2011.07751.x

Mello C V., Clayton DF. 1994. Song-induced ZENK gene expression in auditory pathways of songbird brain and its relation to the song control system. J Neurosci 14:6652–66. doi:0270-6474/94/146652-l 5

Mello C V, Ribeiro S. 1998. ZENK protein regulation by song in the brain of songbirds. J Comp Neurol 393:426–438. doi:10.1002/(SICI)1096-9861(19980420)393:4<426::AID-CNE3>3.0.CO;2-2

Mello C V, Vicario DS, Clayton DF. 1992. Song presentation induces gene expression in the songbird forebrain. Proc Natl Acad Sci U S A 89:6818–22. doi:10.1073/pnas.89.15.6818

Metherato R, Weinberger NM. 1989. Acetylcholine produces stimulus-specific receptive field alterations in cat auditory cortex. Brain Res 480:372–377. doi:10.1016/0006-8993(89)90210-2

Mohebi A, Pettibone JR, Hamid AA, Wong JMT, Vinson LT, Patriarchi T, Tian L, Kennedy RT, Berke JD. 2019. Dissociable dopamine dynamics for learning and motivation. Nature 570:65–70. doi:10.1038/s41586-019-1235-y

Neary JT, Meneely RL, Grever MR, Diven WF. 1972. The interactions between biogenic amines and pyridoxal, pyridoxal phosphate, and pyridoxal kinase. Arch Biochem Biophys 151:42– 47. doi:10.1016/0003-9861(72)90470-5

Nordmann GC, Malkemper EP, Landler L, Ushakova L, Nimpf S, Heinen R, Schuechner S, Ogris E, Keays DA. 2020. A high sensitivity ZENK monoclonal antibody to map neuronal activity in Aves. Sci Rep 10:1–9. doi:10.1038/s41598-020-57757-6

Olesen KM, Auger AP. 2008. Dopaminergic activation of estrogen receptors induces fos expression within restricted regions of the neonatal female rat brain. PLoS One 3. doi:10.1371/journal.pone.0002177

Ono S, Okanoya K, Seki Y. 2016. Hierarchical emergence of sequence sensitivity in the songbird auditory forebrain. J Comp Physiol A 202:163–183. doi:10.1007/s00359-016-1070-7

Pachitariu M, Steinmetz N, Kadir S, Carandini M, Harris KD. 2016. Kilosort: realtime spike- sorting for extracellular electrophysiology with hundreds of channels. bioRxiv. doi:10.1101/061481

Pennartz CMA, Dolleman-Van der Weel MJ, Kitai ST, Lopes Da Silva FH. 1992. Presynaptic dopamine D1 receptors attenuate excitatory and inhibitory limbic inputs to the shell region of the rat nucleus accumbens studied in vitro. J Neurophysiol 67:1325–1334. doi:10.1152/jn.1992.67.5.1325

Phan ML, Pytte CL, Vicario DS. 2006. Early auditory experience generates long-lasting memories that may subserve vocal learning in songbirds. Proc Natl Acad Sci U S A 103:1088–1093. doi:10.1073/pnas.0510136103

Razak KA, Fuzessery ZM. 2009. GABA shapes selectivity for the rate and direction of frequency-modulated sweeps in the auditory cortex. J Neurophysiol 102:1366–1378. doi:10.1152/jn.00334.2009

Reichenbach N, Herrmann U, Kähne T, Schicknick H, Pielot R, Naumann M, Dieterich DC, Gundelfinger ED, Smalla K-H, Tischmeyer W. 2015. Differential effects of dopamine signalling on long-term memory formation and consolidation in rodent brain. Proteome Sci 13:13. doi:10.1186/s12953-015-0069-2

Reiner A, Medina L, Veenman CL. 1998. Structural and functional evolution of the basal ganglia in vertebrates. Brain Res Rev 28:235–285. doi:10.1016/S0165-0173(98)00016-2

Reiner, Karle EJ, Anderson KD, Medina L. 1994. Catecholaminergic perikarya and fibers in the avian nervous system. Phylogeny Dev Catech Syst CNS Vertebr 135–181.

Remage-Healey L, Coleman MJ, Oyama RK, Schlinger B a. 2010. Brain estrogens rapidly strengthen auditory encoding and guide song preference in a songbird. Proc Natl Acad Sci U S A 107:3852–3857. doi:10.1073/pnas.0906572107

Remage-Healey L, Dong SM, Chao a., Schlinger B a. 2012. Sex-specific, rapid neuroestrogen fluctuations and neurophysiological actions in the songbird auditory forebrain. J Neurophysiol 107:1621–1631. doi:10.1152/jn.00749.2011

Remage-Healey L, Joshi NR. 2012. Changing Neuroestrogens Within the Auditory Forebrain Rapidly Transform Stimulus Selectivity in a Downstream Sensorimotor Nucleus. J Neurosci 32:8231–8241. doi:10.1523/JNEUROSCI.1114-12.2012

Remage-Healey L, Maidment NT, Schlinger B a. 2008. Forebrain steroid levels fluctuate rapidly during social interactions. Nat Neurosci 11:1327–1334. doi:10.1038/nn.2200

Rodríguez-Saltos CA, Lyons SM, Sockman KW, Maney DL. 2018. Sound-induced monoaminergic turnover in the auditory forebrain depends on endocrine state in a seasonally-breeding songbird. J Neuroendocrinol e12606. doi:10.1111/jne.12606

Rothman JS, Silver RA. 2018. Neuromatic: An integrated open-source software toolkit for acquisition, analysis and simulation of electrophysiological data. Front Neuroinform 12:1– 21. doi:10.3389/fninf.2018.00014

Saldanha CJ, Schlinger BA, Micevych PE, Horvath TL. 2004. Presynaptic N-Methyl-D-Aspartate Receptor Expression Is Increased by Estrogen in an Aromatase-Rich Area of the Songbird Hippocampus. J Comp Neurol 469:522–534. doi:10.1002/cne.11035

Saldanha CJ, Tuerk MJ, Kim YH, Fernandes AO, Arnold AP, Schlinger BA. 2000. Distribution and regulation of telencephalic aromatase expression in the zebra finch revealed with a specific antibody. J Comp Neurol 423:619–630. doi:10.1002/1096-9861(20000807)423:4<619::AID-CNE7>3.0.CO;2-U

Schicknick H, Reichenbach N, Smalla KH, Scheich H, Gundelfinger ED, Tischmeyer W. 2012. Dopamine modulates memory consolidation of discrimination learning in the auditory cortex. Eur J Neurosci 35:763–774. doi:10.1111/j.1460-9568.2012.07994.x

Schicknick H, Schott BH, Budinger E, Smalla KH, Riedel A, Seidenbecher CI, Scheich H, Gundelfinger ED, Tischmeyer W. 2008. Dopaminergic modulation of auditory cortex-dependent memory consolidation through mTOR. Cereb Cortex 18:2646–2658. doi:10.1093/cercor/bhn026

Schmidt MF, Ding L. 2014. Achieving perfection through variability: the basal ganglia helped me do it! Neuron 82:6–8. doi:10.1016/j.neuron.2014.03.010

Schneider DM, Woolley SMN. 2013. Sparse and Background-Invariant Coding of Vocalizations in Auditory Scenes. Neuron 79:141–152. doi:10.1016/j.neuron.2013.04.038

Scully EN, Hahn AH, Campbell KA, McMillan N, Congdon J V., Sturdy CB. 2017. ZENK expression following conspecific and heterospecific playback in the zebra finch auditory forebrain. Behav Brain Res 331:151–158. doi:10.1016/j.bbr.2017.05.023

Sherman AD, Mott J. 1985. Amphetamine stimulation of glutaminase is blocked by neuroleptics. Life Sci 36:1163–1167. doi:10.1016/0024-3205(85)90233-4

Staley KJ, Longacher M, Bains JS, Yee A. 1998. Presynaptic modulation of CA3 network activity. Nat Neurosci 1:201–209. doi:10.1038/651

Takeuchi T, Duszkiewicz AJ, Sonneborn A, Spooner PA, Yamasaki M, Morris RGM, Watanabe M, Smith CC, Fernández G, Deisseroth K, Robert W. 2016. Locus coeruleus and dopaminergic consolidation of everyday memory. Nature 537:357–362. doi:10.1038/nature19325

Tanaka M, Sun F, Li Y, Mooney R. 2018. A mesocortical dopamine circuit enables the cultural transmission of vocal behaviour. Nature. doi:10.1038/s41586-018-0636-7

Tokarev K, Tiunova A, Scharff C, Anokhin K. 2011. Food for song: Expression of C-Fos and ZENK in the zebra finch song nuclei during food aversion learning. PLoS One 6. doi:10.1371/journal.pone.0021157

Tozzi A, de Iure A, Tantucci M, Durante V, Quiroga-Varela A, GiampÃ C, Di Mauro M, Mazzocchetti P, Costa C, Di Filippo M, Grassi S, Pettorossi VE, Calabresi P. 2015. Endogenous 17b-estradiol is required for activity-dependent long-term potentiation in the striatum: interaction with the dopaminergic system. Front Cell Neurosci 9:192. doi:10.3389/fncel.2015.00192

Tremblay R, Lee S, Rudy B. 2016. GABAergic Interneurons in the Neocortex: From Cellular Properties to Circuits. Neuron 91:260–292. doi:10.1016/j.neuron.2016.06.033

Tsunada J, Lee JH, Cohen YE. 2012. Differential representation of auditory categories between cell classes in primate auditory cortex. J Physiol 590:3129–3139. doi:10.1113/jphysiol.2012.232892

Vahaba DM, Macedo-Lima M, Remage-Healey L. 2017. Sensory Coding and Sensitivity to Local Estrogens Shift during Critical Period Milestones in the Auditory Cortex of Male Songbirds. eNeuro 4:ENEURO.0317-17.2017. doi:10.1523/ENEURO.0317-17.2017

von Eugen K, Tabrik S, Güntürkün O, Ströckens F. 2020. A comparative analysis of the dopaminergic innervation of the executive caudal nidopallium in pigeon, chicken, zebra finch, and carrion crow. J Comp Neurol 1–27. doi:10.1002/cne.24878

Wallhäusser-Franke E, Collins CE, DeVoogd TJ. 1995. Developmental changes in the distribution of NADPH-diaphorase-containing neurons in telencephalic nuclei of the zebra finch song system. J Comp Neurol 356:345–354.

Wang J, McFadden SL, Caspary D, Salvi R. 2002. Gamma-aminobutyric acid circuits shape response properties of auditory cortex neurons. Brain Res 944:219–231. doi:10.1016/S0006-8993(02)02926-8

Wu GK, Arbuckle R, Liu B hua, Tao HW, Zhang LI. 2008. Lateral Sharpening of Cortical Frequency Tuning by Approximately Balanced Inhibition. Neuron 58:132–143. doi:10.1016/j.neuron.2008.01.035

Yague JG, Muñoz A, De Monasterio-Schrader P, DeFelipe J, Garcia-Segura LM, Azcoitia I. 2006. Aromatase expression in the human temporal cortex. Neuroscience 138:389–401. doi:10.1016/j.neuroscience.2005.11.054

Yanagihara S, Ikebuchi M, Mori C, Tachibana RO, Okanoya K. 2019. Social context modulates auditory activity in a songbird VTA/SNcProgram No. 233.03. 2019 Neuroscience Meeting Planner. Chicago, IL.

Yanagihara S, Yazaki-Sugiyama Y. 2016. Auditory experience-dependent cortical circuit shaping for memory formation in bird song learning. Nat Commun 7:11946. doi:10.1038/ncomms11946

Yildiz ZT, Woolley SC. 2017. Singing modulates parvalbumin interneurons throughout songbird forebrain vocal control circuitry. PLoS One 12:1–16. doi:10.1371/journal.pone.0172944

Zhang L, Bose P, Warren RA. 2014. Dopamine Preferentially Inhibits NMDA Receptor-Mediated EPSCs by Acting on Presynaptic D1 Receptors in Nucleus Accumbens during Postnatal Development. PLoS One 9:e86970. doi:10.1371/journal.pone.0086970

